# Ubiquitin-driven protein condensation initiates clathrin-mediated endocytosis

**DOI:** 10.1101/2023.08.21.554139

**Authors:** Feng Yuan, Sadhana Gollapudi, Kasey J. Day, Grant Ashby, Arjun Sangani, Brandon T. Malady, Liping Wang, Eileen M. Lafer, Jon M. Huibregtse, Jeanne C. Stachowiak

## Abstract

Clathrin-mediated endocytosis is an essential cellular pathway that enables signaling and recycling of transmembrane proteins and lipids. During endocytosis, dozens of cytosolic proteins come together at the plasma membrane, assembling into a highly interconnected network that drives endocytic vesicle biogenesis. Recently, multiple groups have reported that early endocytic proteins form flexible condensates, which provide a platform for efficient assembly of endocytic vesicles. Given the importance of this network in the dynamics of endocytosis, how might cells regulate its stability? Many receptors and endocytic proteins are ubiquitylated, while early endocytic proteins such as Eps15 contain ubiquitin-interacting motifs. Therefore, we examined the influence of ubiquitin on the stability of the early endocytic protein network. In vitro, we found that recruitment of small amounts of polyubiquitin dramatically increased the stability of Eps15 condensates, suggesting that ubiquitylation could nucleate endocytic assemblies. In live cell imaging experiments, a version of Eps15 that lacked the ubiquitin-interacting motif failed to rescue defects in endocytic initiation created by Eps15 knockout. Furthermore, fusion of Eps15 to a deubiquitylase enzyme destabilized nascent endocytic sites within minutes. In both in vitro and live cell settings, dynamic exchange of Eps15 proteins, a hallmark of liquid-like systems, was modulated by Eps15-Ub interactions. These results collectively suggest that ubiquitylation drives assembly of the flexible protein network responsible for catalyzing endocytic events. More broadly, this work illustrates a biophysical mechanism by which ubiquitylated transmembrane proteins at the plasma membrane could regulate the efficiency of endocytic recycling.

**Significance Statement:** The assembly of proteins into dynamic, liquid-like condensates is an emerging principle of cellular organization. During clathrin-mediated endocytosis, a liquid-like protein network catalyzes vesicle assembly. How do cells regulate these assemblies? Here we show that ubiquitin and endocytic proteins form a dynamic, mutually-reinforcing protein network in vitro and in live cells. To probe the impact of ubiquitylation on the dynamics of endocytosis, we engineered opto-genetic control over recruitment of proteins to nascent endocytic sites. While recruitment of wildtype proteins promoted endocytosis, recruitment of deubiquitylases, enzymes capable of removing ubiquitin, resulted in disassembly of endocytic sites within minutes. These results illustrate that ubiquitylation can regulate the fate of endocytic structures, elucidating a functional connection between protein condensates, endocytosis, and ubiquitin signaling.

## Introduction

Endocytosis, which is responsible for internalizing proteins and lipids from the plasma membrane, is essential for a myriad of cellular functions including signaling, nutrient import, and recycling (1). In the earliest moments of clathrin-mediated endocytosis, the best understood pathway of cellular internalization (2), initiator proteins including Eps15, Fcho, and Intersectin assemble together to create a nascent endocytic site (2, 3). The resulting network of initiators recruits adaptor proteins, such as AP2, CALM/AP180, and Epsin, among many others, which in turn recruit clathrin triskelia (2, 4). Assembly of triskelia into an icosahedral lattice works in concert with adaptor proteins to induce membrane curvature and vesicle budding (2, 4). Transmembrane cargo proteins are recruited throughout the initiation and growth of endocytic structures. Many cargo proteins contain biochemical motifs that mediate binding to endocytic adaptor proteins (5), while post-translational modifications, such as ubiquitylation, drive uptake of transmembrane proteins destined for degradation or recycling (6–9). Once the clathrin coat is fully assembled and loaded with cargo proteins, scission occurs, resulting in the formation of clathrin-coated vesicles that bud off from the plasma membrane, followed by uncoating (2, 3).

Interestingly, the early initiator proteins of clathrin-mediated endocytosis are not incorporated into endocytic vesicles (10, 11). Instead, these proteins function like catalysts, remaining at the plasma membrane to initiate multiple rounds of endocytosis. To promote growth of a clathrin-coated vesicle, the initiator network must remain flexible, allowing adaptors, cargo proteins, and clathrin triskelia to accumulate and rearrange. Similarly, as the vesicle matures, the network of initiators must ultimately dissociate from the nascent vesicle, allowing it to depart into the cytosol. In line with these requirements, recent work has shown that initiator proteins, which contain substantial regions of intrinsic disorder, form highly flexible, liquid-like assemblies that undergo rapid exchange (12–15). Specifically, Day and colleagues showed that Eps15 and Fcho1/2 form liquid-like droplets in vitro. In live cell imaging experiments, this network exhibits optimal catalytic activity when the level of molecular exchange is maintained within an intermediate range (12). When the network was very weak, corresponding to a disassembled state in vitro, the fraction of endocytic events that were short-lived, likely aborting without creating a vesicle, increased. Conversely, an excessively strong initiator network led to the accumulation of overly stable, stalled endocytic structures, corresponding to a solid-like network in vitro. However, an initiator network of intermediate strength, corresponding to a liquid-like state in vitro, optimized the productivity of endocytosis. Work by Wilfling and colleagues suggests that overly stable, solid-like endocytic sites mature into autophagic structures, rather than resulting in endocytosis (13, 14). More broadly, Kozak and colleagues found that Ede1, the yeast homolog of Eps15, mediates assembly of liquid-like condensates that incorporate many endocytic components (15). Similarly, Drawidge et al recently reported that the AtEH1/2 subunits of the TPLATE complex, which are partially homologous to Eps15/Ede1, are thought to mediate protein condensation during clathrin-mediated endocytosis in plants, suggesting that the requirement for a flexible network of endocytic initiator proteins is broadly conserved (16).

Motivated by these findings, we set out to understand how cells regulate the stability of the early endocytic network. As noted above, Eps15/Ede1 is a key component of this network. Interestingly, several studies have suggested that ubiquitylation can play an important role in mediating interactions between Eps15, cargo proteins, and endocytic adaptor proteins (9, 17–20). Specifically, Ede1 contains a ubiquitin association domain near its C terminus, which enables interactions between Ede1 and ubiquitinated proteins (9, 17). Similarly, two ubiquitin interacting motifs (UIMs) at the C terminus of Eps15 are essential for its interactions with ubiquitinated proteins (19). Further, recent work suggests that deubiquitylases help to avoid stalling of endocytic structures in budding yeast (21).

These findings led us to ask whether ubiquitylation could drive the assembly of liquid-like networks of early endocytic proteins. Using purified proteins in vitro, we found that recruitment of small amounts of polyubiquitin significantly enhanced the stability of liquid-like Eps15 droplets, suggesting a potential role for ubiquitylation in nucleating endocytic sites. In live cell imaging experiments we observed that expression of UIM-deficient Eps15 in cells lacking endogenous Eps15 failed to rescue the defect in clathrin-mediated endocytosis caused by Eps15 knockout. Similarly, removing ubiquitin from Eps15 and its close interactors by recruitment of an Eps15 variant containing a broad-spectrum deubiquitylase domain resulted in a significant destabilization of nascent endocytic sites. Using an optogenetic approach, we found that this destabilization occurred within minutes following recruitment of deubiquitylases to endocytic sites. Tying the in vitro and live cell results together, we found that more rapid molecular exchange of Eps15 at both endocytic sites and in vitro condensates results from loss of interactions between Eps15 and ubiquitin, suggesting that ubiquitylation stabilizes recruitment of Eps15 during endocytosis. Finally, loss of Eps15-ubiquitin interactions produces a marked defect in cellular uptake of transferrin, demonstrating the significance of these interactions for receptor recycling. Collectively, these results suggest that ubiquitylation catalyzes clathrin-mediated endocytosis by driving the assembly of a liquid-like initiator protein network.

## Results

### Polyubiquitin partitions strongly to liquid-like droplets of Eps15

Previous work has shown that Eps15 forms liquid-like condensates in vitro (12). To probe the impact of ubiquitin on condensation of Eps15, we compared the partitioning of monoubiquitin (MonoUb) and lysine-63-linked tetra-ubiquitin (K63 TetraUb) into Eps15 droplets. Notably, K63-linked polyubiquitin chains are thought to play an important role in recycling of receptors from the cell surface (22–25). Eps15 is composed of three major domains: an N-terminal region, which consists of three Eps15 Homology (EH) domains; a central coiled-coil domain, which is responsible for dimerization of Eps15; and an intrinsically disordered C-terminal domain (26). The C-terminal domain contains a binding site for the α-subunit of the clathrin adaptor-protein complex AP2, two ubiquitin-interacting motifs (UIMs) (Figure 1a, top), and 15 tripeptide Asp-Pro-Phe (DPF) motifs (26–28). The DPF motifs mediate oligomerization of Eps15 by binding to its N-terminal EH domains (26–28). As previously reported, full-length Eps15 assembles into liquid-like droplets through these multivalent interaction when added to a solution of 3% w/v PEG8000 in a buffer consisting of 20 mM Tris-HCl, 150 mM NaCl, 5 mM TCEP, 1 mM EDTA and 1 mM EGTA at pH 7.5 (12) (Figure 1a, bottom, where ten percent of Eps15 molecules were labeled with Atto 488 for visualization).

**Figure 1.**
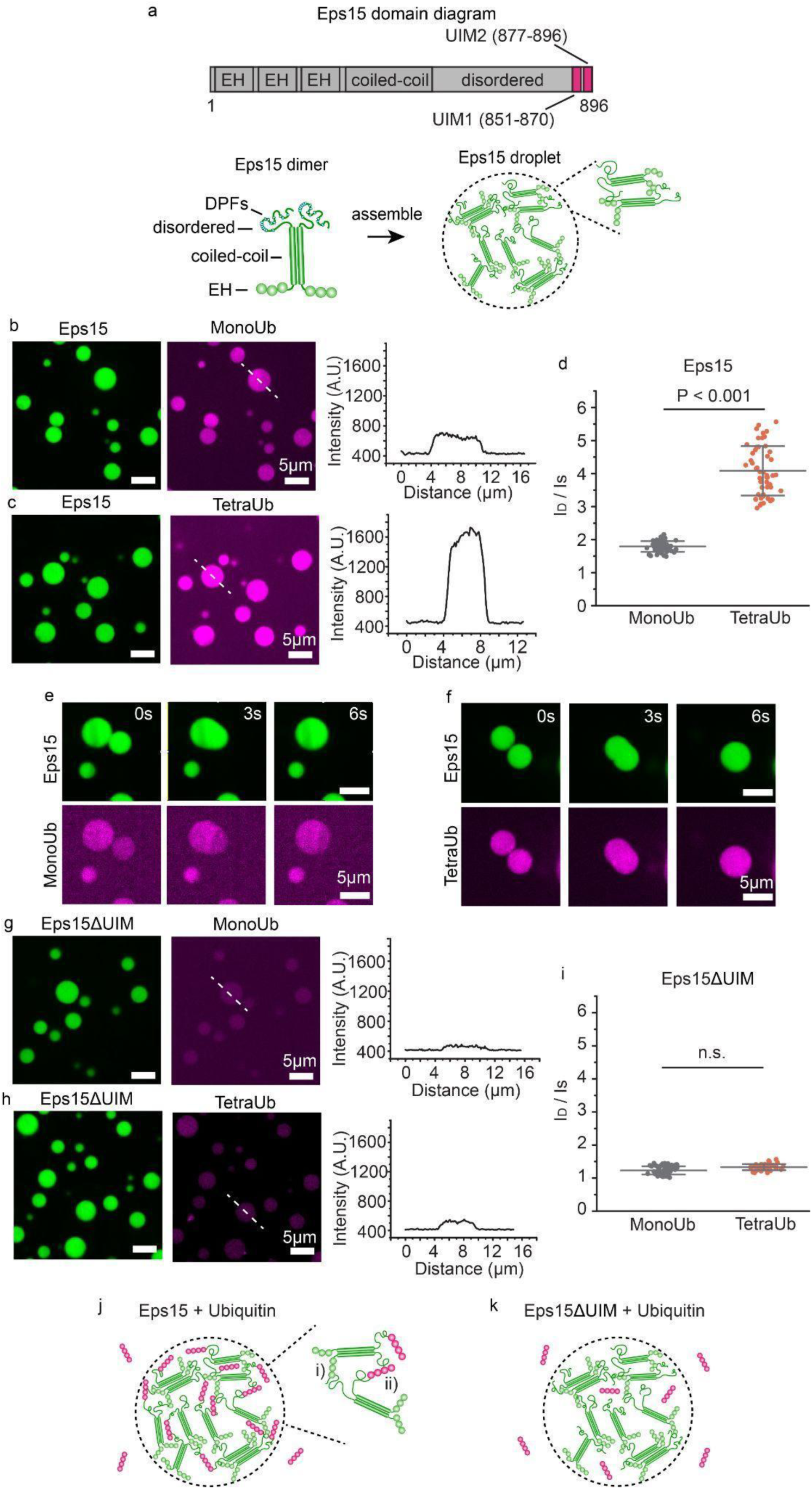
Polyubiquitin partitions strongly into liquid-like droplets of Eps15. **a**, top: Schematic of Eps15 functional domains. Eps15 consists of three EH domains at its N terminus followed by a coiled-coil domain and a long disordered region containing two ubiquitin interacting motifs (UIMs) at the C terminal end. Bottom: cartoons depict domain organization of Eps15 in dimeric form. 15 tripeptide Asp-Pro-Phe (DPF) motifs are interspersed throughout the disordered region, which can bind the EH domains and allow itself to assemble into liquid-like droplets. **b, c,** Eps15 (7μM) droplets (green) incubated with 1μM MonoUb and 0.25μM K63 linkage TetraUb (magenta), respectively. Plots on the right depict intensity profile of ubiquitin channel along the white dashed line shown in the corresponding images. **d**, The distribution of the ubiquitin intensity ratio between the intensity inside the droplets (I_D_) and the solution (I_S_). In total 50 droplets were analyzed under each condition. **e, f**, Representative time course of fusion events between droplets containing Eps15 and MonoUb (**e**) and droplets containing Eps15 and TetraUb (**f**). **g-i**, Same with **b-d** except that droplets were formed with Eps15 mutant, Eps15ΔUIM, with the depletion of the two UIMs (aa 851-896). **j, k,** Pictorial representation of ubiquitin binding and partitioning into Eps15 droplets through interaction with UIMs at the C terminus of Eps15 (**j**) and deletion of UIMs impairs ubiquitin partitioning into Eps15 droplets (**k**). Inset in **j** shows the two types of interactions in Eps15-polyubiquitin network: i) DPF motif interacting with EH domain, and ii) polyubiquitin interacting with UIM domains. All droplet experiments were performed in 20 mM Tris-HCl, 150 mM NaCl, 5 mM TCEP, 1 mM EDTA and 1 mM EGTA at pH 7.5 with 3% w/v PEG8000. Error bars are standard deviation. Statistical significance was tested using an unpaired, two-tailed student’s t test. All scale bars equal 5 μm.

To examine the impact of ubiquitin on phase separation of Eps15, MonoUb and TetraUb, labeled with Atto 647, were added to Eps15 droplets at a final concentration of 1 μM and 0.25 μM, respectively, maintaining an equivalent mass per volume of ubiquitin. Images of the Eps15 droplets were collected after a 5-minute incubation. As shown in Figure 1b, c, both MonoUb and TetraUb partitioned uniformly into Eps15 droplets. However, the partition coefficient of TetraUb into Eps15 droplets, indicated by the ratio between the intensity of ubiquitin in the droplet (I_D_) and the surrounding solution (I_S_), was about twice that of MonoUb (4.1 ± 0.8 vs. 1.8 ± 0.2, Figure 1d), suggesting that Eps15 interacted more strongly with TetraUb. Importantly, Eps15 droplets incubated with either MonoUb or TetraUb remained liquid-like, readily fusing and re-rounding upon contact (Figure 1e, f and Supplementary Information movie S1 and S2). MonoUb and TetraUb partitioned much more weakly into droplets consisting of a version of Eps15 that lacked its UIMs, Eps15ΔUIM (Δaa 851-896), (1.4 ± 0.1 and 1.5 ± 0.1 respectively, with no significant difference, Figure 1g, h and i). We observed a similar phenomenon when Eps15 proteins bound to the surfaces of giant unilamellar vesicles, Supplementary Figure S1. Here addition of TetraUb to membrane vesicles coated with Eps15 resulted in assembly of membrane-bound condensates (Figure S1 a-d, g, h). This effect required Eps15’s UIM domain and was absent when MonoUb was added rather than TetraUb (Figure S1, e, f, i, j). Collectively, these results confirm that Eps15’s UIMs drive the partitioning of ubiquitin into Eps15 droplets, where both Eps15-Eps15 interactions and Eps15-ubiquitin interactions are expected to exist within droplets (Figure 1j, k).

### Loss of Eps15’s ubiquitin interacting motifs eliminates the protein’s role in endocytic initiation

Having demonstrated that multivalent interactions with ubiquitin can promote condensation of Eps15 networks *in vitro*, we next sought to evaluate the impact of interactions between Eps15 and ubiquitin on the dynamics of endocytic events in cells. As discussed in the introduction, previous work has demonstrated that Eps15 plays an important role in stabilizing complexes of early endocytic proteins (29). Importantly, the dynamic assembly of endocytic structures at the plasma membrane is a highly stochastic process. While most assemblies of endocytic proteins ultimately result in productive endocytic events, a minority are unstable, failing to develop into endocytic vesicles. In general, endocytic assemblies that persist at the plasma membrane for less than 20 seconds are regarded as “short-lived” structures (30). These unstable assemblies, which typically consist of a small number of proteins, are thought to spontaneously disassemble rather than forming productive vesicles. In contrast, productive assemblies typically form vesicles and depart from the plasma membranes over timescales of 20 seconds to several minutes. Structures that persist at the plasma membrane for longer periods are characterized as “long-lived” (31). These overly stable assemblies, which are often larger than productive endocytic structures, may fail to develop into endocytic vesicles and could be removed from the plasma membrane by autophagy (13, 14). Shifts in the lifetime of endocytic structures provide insight into the overall efficiency of endocytosis. While an increase in short-lived structures indicates a reduction in stability, an increase in productive or long-lived structures indicates an increase in stability.

We evaluated endocytic dynamics in a human breast cancer-derived epithelial cell line, SUM159, which was gene edited to (i) knockout Eps15, and (ii) add a C-terminal halo-tag to the sigma2 subunit of AP2. As AP2 is the major adaptor protein of the clathrin-mediated endocytic pathway (32, 33), the halo tag, bound to JF646 dye, was used to visualize and track endocytic structures in real-time during live cell imaging (34). Notably, labeling of AP2 sigma, rather than clathrin itself, is a popular approach for tracking the growth of clathrin-coated structures because AP2 is found exclusively at the plasma membrane (35, 36). In contrast, clathrin is found at the plasma membrane, endosomal membranes, and the trans-Golgi network, creating the potential for imaging artifacts (37, 38).

Imaging experiments were performed using total internal reflection fluorescence (TIRF) microscopy to isolate the plasma membrane of adherent cells. In TIRF images, endocytic structures appear as diffraction-limited fluorescent puncta in the JF646 channel, Figure 2a. Eps15, and its variants, described below, were tagged at their C-termini with mCherry for visualization, and co-localized with puncta of AP2 (JF646), Figure 2a. Based on the literature cited above, we loosely classified endocytic structures that persisted at the plasma membrane for less than 20s as “short-lived”, structures that persisted from 20 – 180 seconds as “productive”, and structures that persisted for longer than 180 seconds as “long-lived”. Endocytic structures within each of these categories were observed in our experiments (Figure 2a) and the distribution of lifetimes for the full population of endocytic structures is shown in Figure 2b. In agreement with a recent report (12), Eps15 knockout cells (Eps15KO) had nearly twice as many short-lived structures compared to wildtype cells with endogenous expression of Eps15 (42.1 ± 2.1% vs. 23.2 ± 3.6%, Figure 2b, c). This defect was effectively rescued by transient expression of wildtype Eps15 in Eps15KO cells (23.7 ± 2.6% vs. 23.2 ± 3.6% short-lived structures, Figure 2b, c) at a modest level that was held constant across all studies of Eps15 mutants described below, Figure S2.

**Figure 2.**
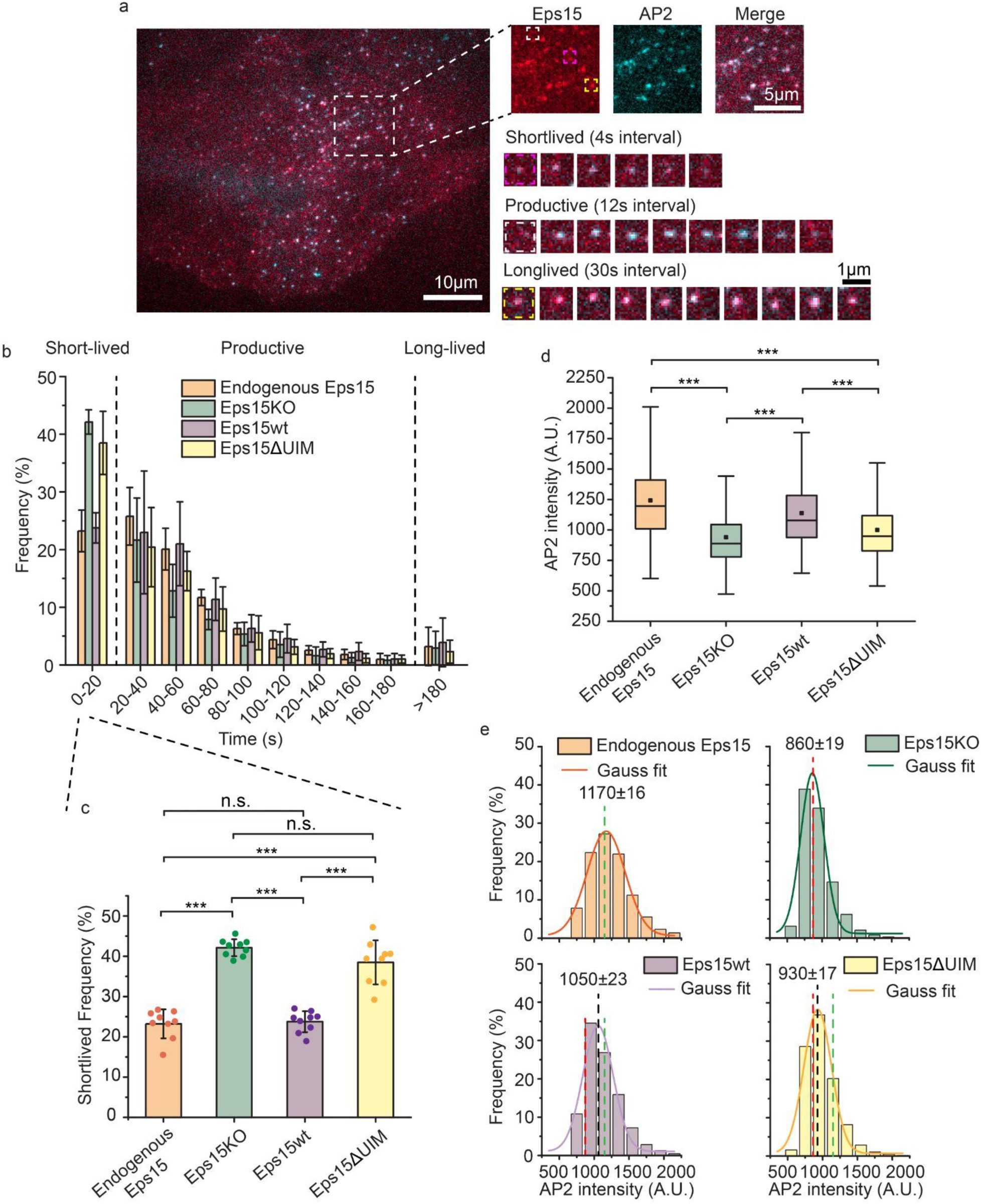
Eps15 knockout creates a significant defect in coated pit dynamics that cannot be rescued by a version of Eps15 that lacks the ubiquitin interacting motif. **a**, Representative image of a SUM cell expressing gene-edited AP-2 σ2-HaloTag: JF646 (cyan) and Eps15-mCherry (red). Large inset highlights three representative clathrin-coated structures shown in smaller insets: short-lived (pink), productive (white) and long-lived (yellow) structures lasting 20 s, 96 s and > 5 min, respectively. Scale bars are labeled in the images. **b**, Histograms of lifetime distributions of clathrin-coated structures under different experimental groups. Endogenous Eps15 represents SUM cells that have endogenous Eps15 expression. Lifetime shorter than 20 s is considered short-lived, lifetime between 20 and 180 s is labeled as productive and structures lasting longer than 180 s are long-lived. Eps15KO represents SUM cells that were CRISPR modified to knockout alleles of endogenous Eps15. Eps15wt and Eps15ΔUIM represent Eps15KO cells transfected with wildtype Eps15 and Eps15 with the depletion of both UIM domains, respectively. mCherry was fused to the C terminus of Eps15 and Eps15ΔUIM for visualization. **c,** bar chart of the short-lived fraction for each group from **b,** error bars are standard deviation, dots represent the results from different cells. **d**, Box plot of endocytic pits AP2 intensity in all four groups. **e**, Histograms and the Gaussian fit of the AP2 intensity distribution tracked in endocytic pits under different experimental groups. Green dotted line indicates the peak distribution in Endogenous Eps15 cells, and red dotted line indicates the peak distribution in Eps15KO cells. For the Endogenous Eps15 group, n = 9 biologically independent cell samples were collected and in total 4346 pits were analyzed. For Eps15KO, n = 9 and 8195 pits. Eps15wt, n = 9 and 4387 pits and Eps15ΔUIM, n = 9, 6779 pits. An unpaired, two-tailed student’s t test was used for statistical significance. n.s. means no significant difference. ***: P < 0.001. All cell images were collected at 37°C.

We also analyzed the impact of Eps15 knockout on the intensity of endocytic structures in the AP2 channel, which, owing to the near 1:1 stoichiometric ratio between AP2 and clathrin (39), serves as a proxy for the maturity and size of endocytic structures. This analysis revealed that Eps15 knockout resulted in significantly reduced AP2 intensity, suggesting less developed endocytic structures, compared to wildtype cells and knockout cells transiently expressing wildtype Eps15, Figure 2d and Figure S3. Specifically, histograms of AP2 intensity showed a shift towards smaller values when Eps15 was knocked out (1170 ± 16, green dotted line, vs. 860 ± 19, red dotted line, in wildtype cells, Figure 2e), a defect which was rescued by transient expression of wildtype Eps15 in knockout cells (1050 ± 23, black dotted line, vs. green dotted line, Figure 2e). Importantly, the very similar distribution of intensities between cells expressing Eps15 endogenously and cells in which Eps15 knockout has been rescued by transient expression of wildtype Eps15 demonstrates that transient expression, at the levels used, does not shift the sizes of endocytic structures or cause structures to fuse together, possible artifacts of over expression. Further, as described under methods, occasional large, non-diffraction-limited structures are excluded from our analysis of endocytic dynamics, reducing sensitivity to potential artifacts. Collectively, these results demonstrate that Eps15 knockout destabilizes endocytic structures, limiting their maturation.

Having established these controls, we next examined the impact of Eps15’s UIMs on Eps15’s ability to promote efficient endocytosis. Specifically, we measured the extent to which a version of Eps15 lacking the UIMs (Eps15ΔUIM, as described above) could rescue the defects created by knockout of the wildtype protein. Interestingly, we found that the elevated number of short-lived endocytic structures observed upon Eps15 knockout was not substantially reduced by expression of Eps15ΔUIM at equivalent levels to the level of Eps15wt required for full rescue, (38.5 ± 5.5% vs. 42.1 ± 2.1%, Figure 2b, c). Similarly, expression of Eps15ΔUIM failed to elevate the average AP2 intensity at endocytic sites above levels measured in knockout cells (Figure 2d, e, black dotted line vs. red line), indicating an inability to stabilize endocytic structures. These results suggest that Eps15’s UIMs are essential to its ability to promote endocytosis.

### Polyubiquitin elevates the melting temperature of liquid-like Eps15 networks

The shorter lifetime of endocytic structures formed when Eps15ΔUIM replaces wild-type Eps15 suggests that loss of the UIMs destabilizes the network of early endocytic proteins. To test this idea, we sought to assess the impact of ubiquitin on the thermodynamic stability of liquid-like condensates of Eps15. For this purpose, we returned to the *in vitro* droplet system in Figure 1 and measured the temperature above which Eps15 droplets dissolved or melted, which is a key indicator of their stability (12). Specifically, when we heated solutions of Eps15, liquid-like Eps15 droplets gradually dissolved, eventually melting such that the solution became homogenous (fully dissolved, Figure 3a-e). The higher the melting temperature, the more energy is needed to prevent proteins from condensing, suggesting a more stable protein network.

**Figure 3.**
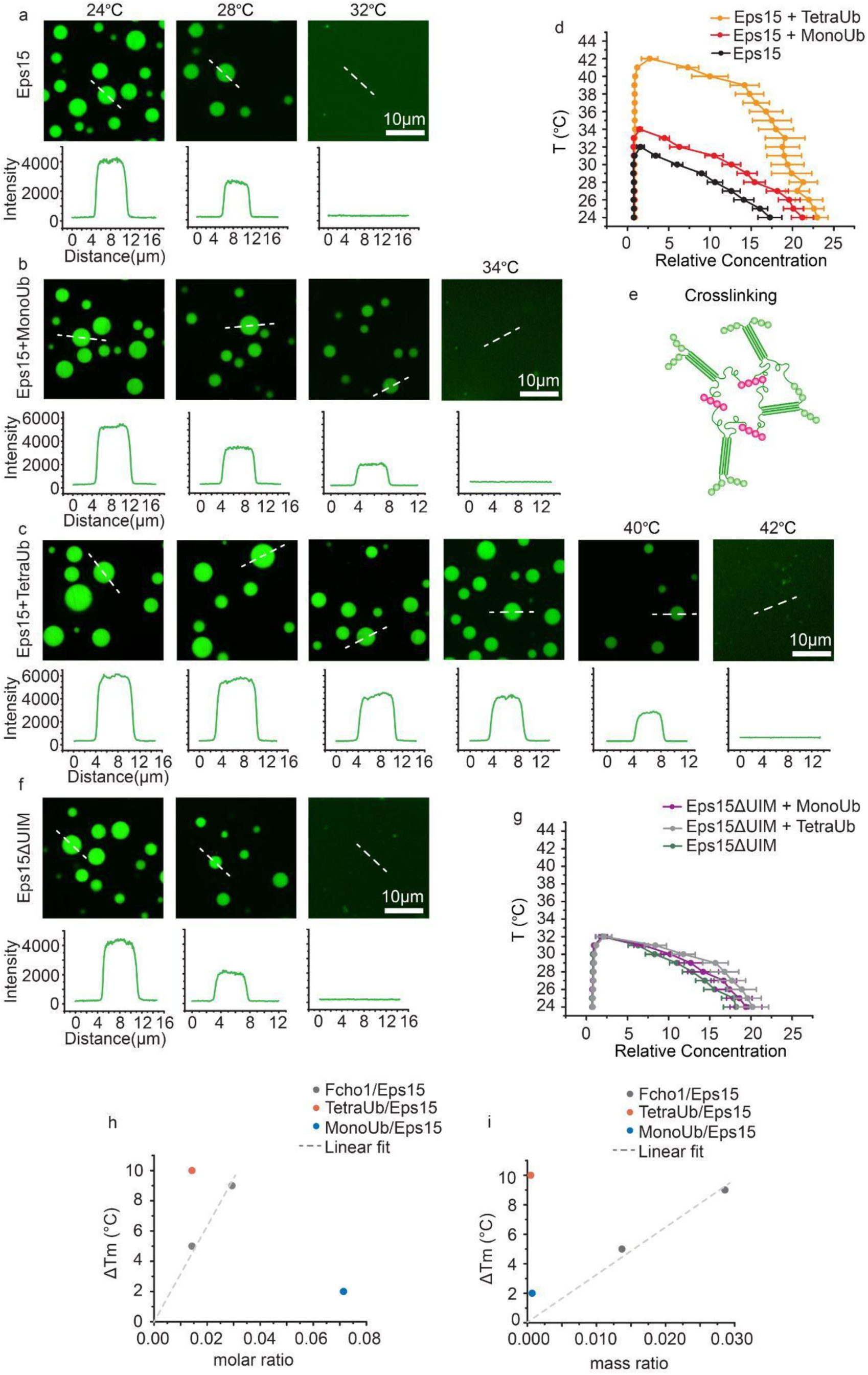
Polyubiquitin elevates the melting temperature of liquid-like Eps15 networks. **a-c, f**, Representative images of protein droplets at increasing temperatures. Plots show fluorescence intensity of Eps15 measured along dotted lines in each image. Droplets are formed from (**a**) 7 μM Eps15, (**b**) 0.5 μM MonoUb, 7 μM Eps15, (**c**) 0.1 μM TetraUb, 7 μM Eps15 and (**f**) 7 μM Eps15ΔUIM in 20 mM Tris-HCl, 150 mM NaCl, 5 mM TCEP, 1 mM EDTA and 1 mM EGTA at pH 7.5 buffer with 3% PEG8000. **d, g**, Phase diagram of Eps15/monoUb/TetraUb (**d**) and Eps15ΔUIM/MonoUb/TetraUb (**g**) droplets mapped by Atto488-labelled Eps15/Eps15ΔUIM fluorescence intensity. Intensity was normalized based on the intensity of the solution. Dots on the right side are protein concentrations in droplets and dots on the left side are concentrations in solution. At least 20 droplets are analyzed under each temperature. Data are mean ± SD. Scale bars equal 10 μm. **e,** Schematic of polyubiquitin crosslinking and stablizing Eps15 network. **h, i**, the change in the melting temperature of Eps15 droplets upon the addition of MonoUb (blue dot), TetraUb (red dot) and Fcho1 (gray dots) of different molar ratio (h) and mass ratio (i). Fcho data were adopted from previous work (12), dash line is the linear fit of Fcho data points.

As shown in Figure 3a, Eps15 droplets (7 μM) gradually dissolved with increasing temperature and melted at approximately 32°C (Figure 3a), in agreement with a previous report (12). Addition of 500 nM MonoUb slightly increased the melting temperature to 34°C (Figure 3b). In contrast, addition of 100 nM TetraUb raised the melting temperature more substantially to 42°C (Figure 3c), suggesting that TetraUb is more effective in stabilizing Eps15 condensates, in comparison to MonoUb. Using these data, we mapped a temperature-concentration phase diagram for Eps15 condensates (Figure 3d). The relative fluorescence intensity of the droplets compared with the surrounding solution provided an estimate of the relative protein concentration in the two phases, C_D_ and C_S_, respectively. These concentrations represent the ends of a tie-line on a temperature-concentration phase diagram at each temperature. As the temperature increased, the intensity of the Eps15 droplets decreased (Figure 3a-c) and the tie-lines became shorter as C_D_ and C_S_ became more similar, Figure 3d. The two concentrations ultimately became equivalent above the melting temperature, owing to dissolution of the droplets, Figure 3d. These results are in line with a recent report showing that poly-ubiquitin can enhance phase separation of proteins involved in protein degradation and autophagy (40).

To assess the impact of Eps15’s UIM domains, we mapped the phase diagram of Eps15ΔUIM droplets in the presence of either MonoUb or TetraUb (Figure 3f, g and Figure S4), keeping protein concentrations the same as those used in experiments with wild-type Eps15. The phase diagrams indicated that neither MonoUb nor TetraUb had a significant impact on the melting temperature of condensates composed of Eps15ΔUIM, demonstrating that the stabilization effect observed with wildtype Eps15 arises from specific interactions between Eps15 and ubiquitin. Collectively, these results suggest that polyubiquitin not only partitions preferentially into Eps15 condensates (Figure 1) but also reinforces the protein network in a UIM-dependent manner (Figure 3e).

Previous work showed that Fcho1 can similarly increase the stability of the Eps15 protein network (12). Interestingly, our experiments demonstrate that the impact of TetraUb is approximately 2-20 fold larger than that of Fcho1, depending on whether equal molar or mass ratios are compared (Figure 3h, i, respectively). This observation highlights the potential for relatively small amounts of polyubiquitin, or multiple interactions with monoubiquitin, either of which could be conjugated to the endocytic machinery or to the intracellular domains of cargo proteins, to drive condensation of the Eps15 network, a key early step in endocytosis.

### Deubiquitination inhibits recruitment of Eps15, destabilizing nascent endocytic structures

If polyubiquitin stabilizes endocytic protein networks, then stripping ubiquitin from the proteins that make up an endocytic site should disrupt the dynamics of endocytosis. To test this prediction, we added a deubiquitylating enzyme (DUB) to Eps15 to remove ubiquitin modifications from Eps15 as well as its close interactors in our live cell imaging experiments. These interactors likely include other endocytic proteins and the intracellular domains of receptors that are internalized by endocytic sites (41–43). Using this approach, we sought to probe the broader sensitivity of endocytic assemblies to loss of ubiquitylation. Here we employed a “broad-spectrum” deubiquitylase, UL36, which consists of the N-terminal domain (residues 15-260) of the type I Herpes virus VP1/2 tegument protein (41, 44). This domain cleaves both K63 and K48-linked polyubiquitin chains (42, 44, 45). K63-linked chains are more traditionally associated with endocytic recycling (24, 46), while K48-linked chains are thought to be mainly involved in targeting proteins for proteasomal degradation (46). UL36 was inserted at the C-terminus of Eps15, prior to the mCherry tag, to create the fusion protein Eps15-DUB (Figure 4a). As a control for the non-specific impact of the DUB fusion on endocytic function, a catalytically inactive UL36, which contains a mutation of its core catalytic residues (Cys65 to Ala) (42, 45), was used to create the chimera, Eps15-DUB-dead (Figure 4a). We then expressed Eps15-DUB and Eps15-DUB-dead in separate populations of Eps15 knockout cells and tracked endocytic dynamics with TIRF microscopy, as described above (Figure 2). TIRF images revealed clear colocalization of Eps15-DUB and Eps15-DUB-dead (mCherry) with endocytic sites, represented by fluorescent puncta in the AP2 (JF646) channel, Figure 4b. Interestingly, Eps15-DUB led to a further decrease in AP2 intensity (Figure 4c and Figure S5, top, 740 ± 9 compared to 860 ± 19) at endocytic sites compared to Eps15 knockout cells alone (Figure S5, red dotted line), suggesting smaller, less mature endocytic sites. In contrast, expression of Eps15-DUB-dead restored AP2 intensity to a level similar to wildtype cells (Figure 4c, and Figure S5, bottom, 1000 ± 21, vs. the green dotted line). Furthermore, the fraction of unstable, short-lived endocytic sites increased by more than 30% in cells expressing Eps15-DUB, compared to cells lacking Eps15 (55.6 ± 4.5% vs. 42.1 ± 2.1% Figure 4d). These results suggest that Eps15-DUB not only failed to rescue the defect caused by Eps15 knockout, but made the defect larger, shifting the balance toward unproductive, short-lived endocytic events. In contrast, Eps15-DUB-dead provided a partial rescue of endocytic dynamics, reducing the fraction of short-lived structures to 33.6 ± 2.2%, Figure 4d. If Eps15 were the only endocytic protein that relied on ubiquitin for its assembly, we would have expected Eps15-DUB to create no greater effect on endocytic dynamics than Eps15ΔUIM. Therefore, our observation of a larger defect suggests that the importance of ubiquitylation to endocytosis extends beyond Eps15, potentially involving other UIM-containing endocytic proteins such as Epsin (17, 19, 47), which could interact with ubiquitin modifications on virtually any endocytic protein or transmembrane cargo protein.

**Figure 4.**
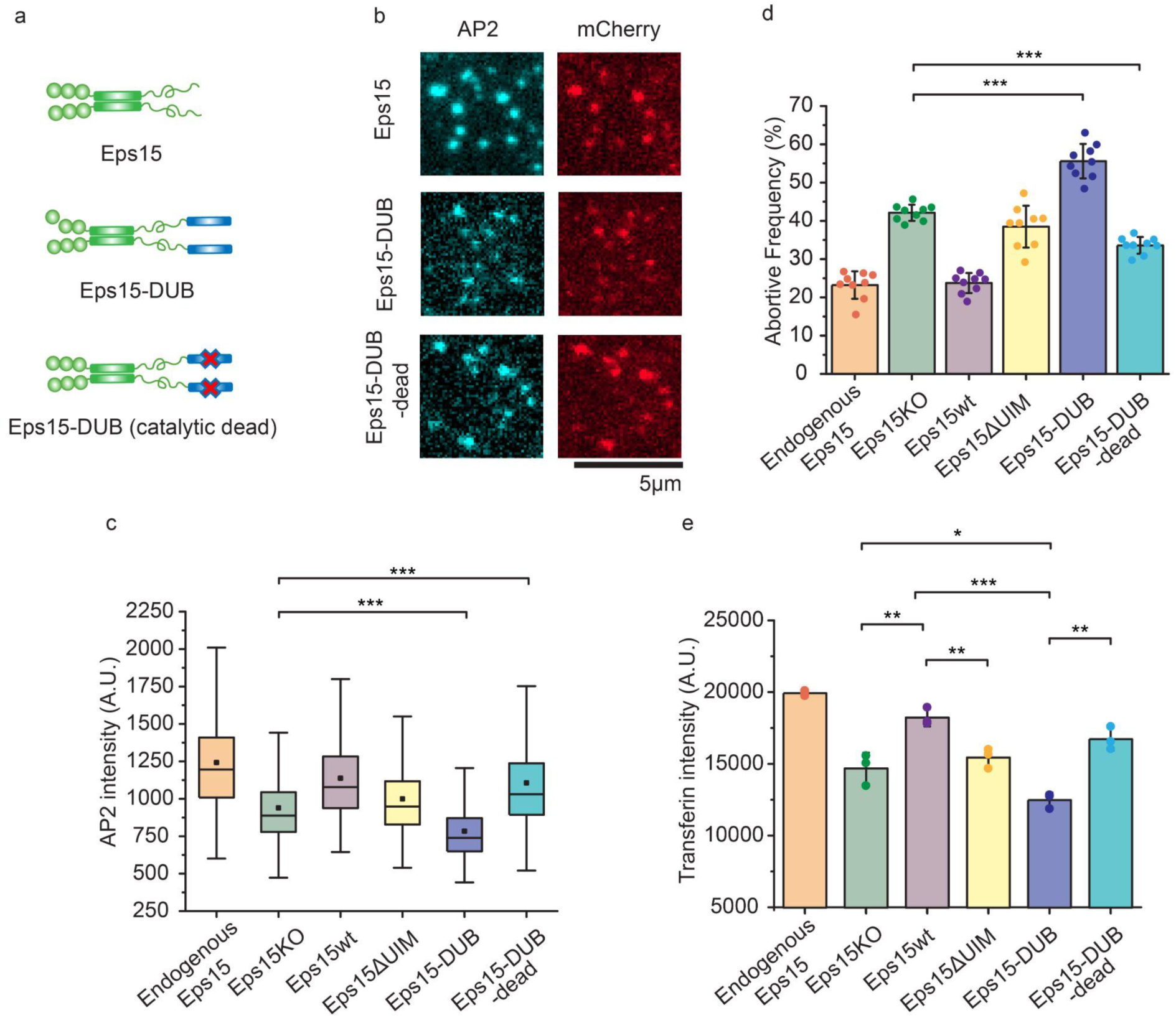
Fusion of a deubiquitinating enzyme to Eps15 results in even more short-lived endocytic events than deletion of Eps15. **a**, Schematic of Eps15 dimeric form, Eps15-DUB (deubiquitylase fused to C terminal end of Eps15), and Eps15-DUB-dead (same with DUB but with a mutation that makes DUB catalytically dead). mCherry is not shown in the cartoon but all three constructs have mCherry at their C terminus for visualization. **b**, Representative images showing Eps15, Eps15-DUB and Eps15-DUB-dead colocalization with AP2 in endocytic sites when Eps15KO SUM cells were transfected to express corresponding proteins. Scale bar = 5 μm. **c, d**, Box plot of endocytic pits AP2 intensity (**c**), and frequency of short-lived structures (**d**) under each condition. Endogenous Eps15, Eps15KO, Eps15wt and Eps15ΔUIM are adopted from the same data shown in Figure 2. For Eps15-DUB, n = 9 biologically independent cell samples were collected and in total 8640 pits were analyzed. For Eps15-DUB-dead, n = 9 and 7420 pits were analyzed. Dots represent the frequency from each sample. **e**, Transferrin uptake under each condition measured by flow cytometry. N = 3 independent samples were measured for each group. An unpaired, two-tailed student’s t test was used for statistical significance. *: P < 0.05, **: P < 0.01, ***: P < 0.001. Error bars represent standard deviation. Cells were imaged at 37°C for all conditions.

### Loss of Eps15-ubiquitin interactions substantially reduces endocytic uptake of transferrin

To what extent do the observed defects in the growth and dynamics of endocytic structures impact the physiological function of endocytosis, which is to recycle transmembrane cargos and their ligands from the cell surface? To address this question, we measured the impact of perturbations to Eps15-ubiquitin interactions on cellular uptake of transferrin, the natural ligand of the transferrin receptor. Transferrin receptor is one of the best-characterized cargo proteins of the clathrin-mediated endocytic pathway, and measurement of transferrin uptake is a classic assay for assessing changes in the efficiency of endocytosis owing to genetic perturbation (48–50). Using flow cytometry, we quantified relative endocytic uptake of fluorescent-labeled transferrin (Atto488), via endocytosis of endogenous transferrin receptors. Non-internalized transferrin bound to the cell surface was removed prior to the measurements as described in the methods section. We examined the same experimental groups used in Figures 4c,d: (i) wildtype cells expressing endogenous Eps15, (ii) Eps15 knockout cells (Eps15KO), (ii) Eps15 knockout cells expressing wildtype Eps15-mCherry (Eps15wt), (iv) Eps15 knockout cells Eps15ΔUIM, (v) Eps15 knockout cells expressing Eps15-DUB, and (vi) Eps15 knockout cells expressing Eps15-DUB-dead, Figure 4e, Figure S6 and S7. Comparing these data to the relative AP2 intensity of endocytic structures (Figure 4c) and the fraction of abortive events (Figure 4d) across the same groups reveals a striking correlation. In each case, perturbations that inhibit endocytosis (Eps15KO, Eps15ΔUIM, Eps15-DUB) drive an increase in the frequency of abortive structures, which consistently correlates with a decrease in relative structure size and transferrin uptake. In all three assays, Eps15 knockdown creates a defect that is fully rescued by re-expression of Eps15wt but which Eps15ΔUIM almost entirely fails to rescue. Further, in each assay, replacement of Eps15wt by Eps15-DUB not only fails to rescue the defect, but increases its magnitude. These results illustrate that changes in endocytic dynamics and growth associated with loss of Eps15-ubiquitin interactions result in corresponding changes in the ability of cells to internalize ligand-bound receptors.

### In condensates and at endocytic sites, loss of Eps15-ubiquitin interactions results in increased molecular exchange of Eps15

Our results so far suggest that loss of interactions with ubiquitin destabilizes Eps15 condensates in vitro and disrupts the assembly of endocytic protein networks in which Eps15 participates. Both observations could be explained by unstable assembly of Eps15 networks in the absence of ubiquitin. Therefore, to compare the stability of Eps15 networks in vitro and in live cells, we performed fluorescence recovery after photobleaching (FRAP) experiments in both contexts. In vitro, we performed FRAP experiments on droplets consisting of wild type Eps15 and Eps15ΔUIM in the presence and absence of MonoUb and TetraUb. These experiments revealed that the rate and extent of Eps15 exchange inside protein condensates decreased 2.6-fold when TetraUb was introduced (1.4-fold for MonoUb), reductions which did not occur when droplets consisted of Eps15ΔUIM (Figure 5a-c). Similarly, to examine the rate of Eps15 exchange in live cells, we measured FRAP of mCherry-labeled Eps15 variants at endocytic sites (Figure 5d,e). While wild type Eps15 displayed partial (41.3%) recovery with a t_1/2_ of 19 s, Eps15ΔUIM recovered more rapidly and completely (t_1/2_ = 6 s, 81.5% recovery), suggesting less stable recruitment to endocytic sites. Eps15-DUB recovered even more quickly and completely (t_1/2_ = 8 s, 96.7% recovery), suggesting that the loss of ubiquitination at endocytic sites prevents stable recruitment of Eps15. In contrast, Eps15-DUB-dead recovered similarly to the wildtype protein (t_1/2_ = 12s, 46.3% recovery). Interestingly, these changes in stability were mirrored by AP2, which is downstream in the clathrin pathway relative to Eps15 and is required for the growth of endocytic structures, Figure 5f,g. Specifically, unstable recruitment of Eps15, due to removal of the UIM domain or addition of a DUB domain, resulted in unstable recruitment of AP2, helping to explain the downstream impact of these interactions on the physiology of endocytosis, as demonstrated in Figure 4. Collectively, these results demonstrate that Eps15 exchanges rapidly during the growth of endocytic structures, suggesting that it is part of a dynamic network. Further, the similar impact of Eps15-ubiquitin interactions on the exchange of Eps15 in vitro and at endocytic sites suggests that a highly flexible, ubiquitin-dependent network likely exists in both contexts, and that in vitro assays on protein condensates provide a useful tool for probing such interactions. Specifically, when ubiquitin is present at endocytic sites, the Eps15 network is stabilized, driving endocytosis productively forward. In contrast, when ubiquitin is absent, the Eps15 network is destabilized, increasing the probability of abortive endocytic events, Figure 5h.

**Figure 5.**
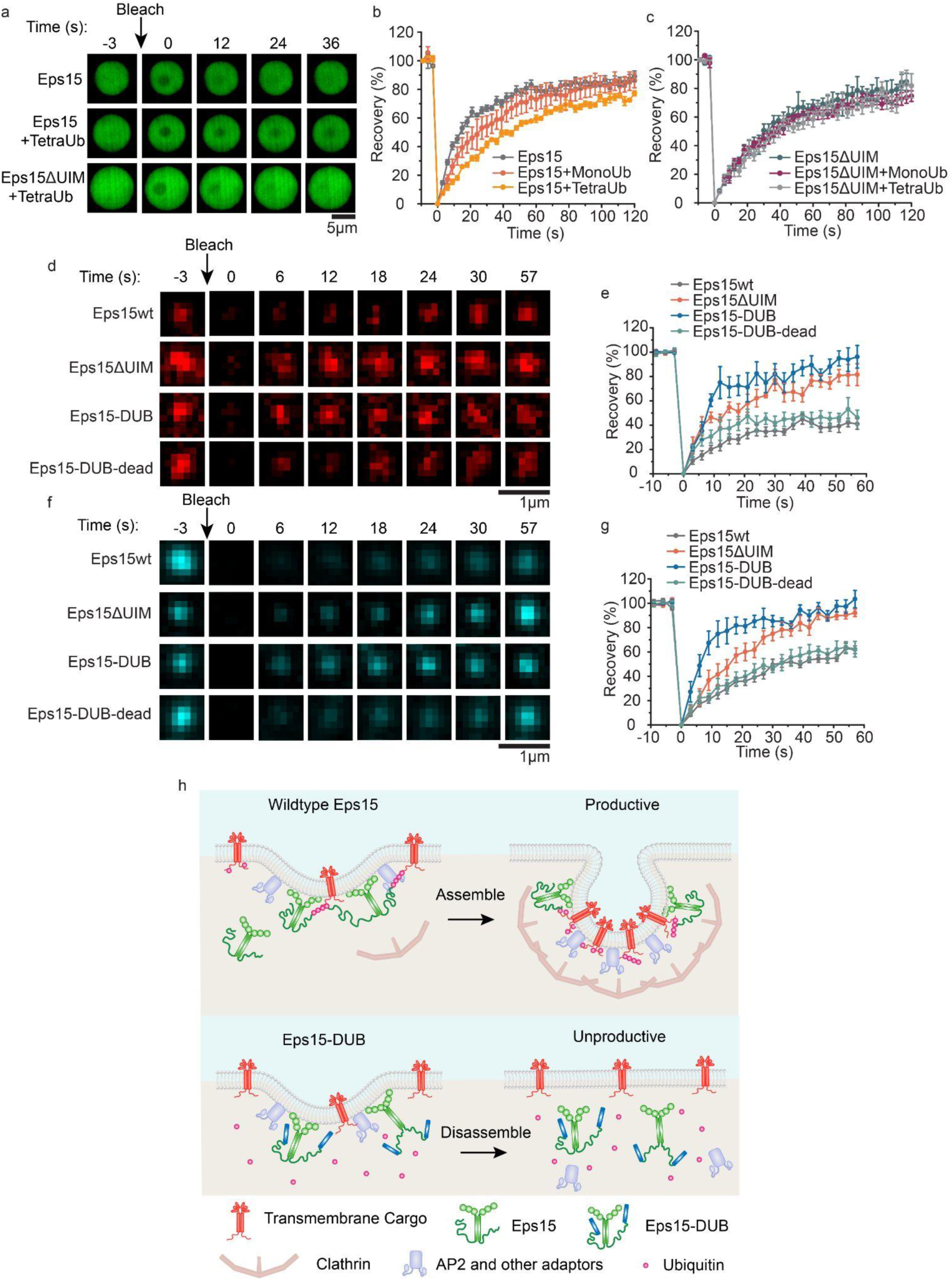
In condensates and at endocytic sites, loss of Eps15-ubiquitin interactions results in increased molecular exchange of Eps15. **a**, Representative mage series of fluorescence recovery after photobleaching of Eps15, Eps15 + TetraUb, and Eps15ΔUIM + TetraUb droplets, respectively. Scale bar = 5 μm. **b, c**, Fluorescence recovery curves for Eps15 (**b**) or Eps15ΔUIM (**c**) droplets in the presence of MonoUb or TetraUb. Eps15/Eps15ΔUIM concentration was maintained at 7 μM with the addition of 1 μM MonoUb and 0.25 μM TetraUb, respectively. Droplets were made in 20 mM Tris-HCl, 150 mM NaCl, 5 mM TCEP, 1 mM EDTA and 1 mM EGTA at pH 7.5 with 3% w/v PEG8000 and all droplet experiments were conducted at room temperature. Data shown as mean ± standard error. n = 6 droplets under each condition. Mobile fractions and t_1/2_: Eps15 (85.9 ± 5.3%, 12.5 ± 0.8s), Eps15 + MonoUb (82.7 ± 10.9%, 18.0 ± 1.3s), Eps15 + TetraUb (79.0 ± 9.3%, 31.1 ±1.2s), Eps15ΔUIM (81.5 ±12.1%, 21.4 ± 1.1s), Eps15ΔUIM + MonoUb (75.74 ±6.5%, 21.8 ±1.9s), Eps15ΔUIM + TetraUb (76.3 ± 11.3%, 24.12 ± 1.6s). **d, e**, Representative images of fluorescence recovery of Eps15-mCherry variants in clathrin-coated structures in Eps15 knockout cells expressing corresponding variants (**d**) and the average fluorescence recovery plots for each condition (**e**). Mobile fractions and t_1/2_: Eps15wt (41.3%, 19.1s), Eps15ΔUIM (81.5%, 5.5s), Eps15-DUB (96.7%, 8.0s) and Eps15-DUB-dead (46.3%, 11.5s). **f, g**, Representative images of fluorescence recovery of AP2-HaloTag labeled with JF646 in clathrin-coated structures in Eps15 knockout cells expressing corresponding Eps15 variants (**f**) and the average fluorescence recovery plots (**g**). Mobile fractions and t_1/2_: Eps15wt (71.9%, 29.7s), Eps15ΔUIM (100.0%, 15.1s), Eps15-DUB (103.9%, 6.8s) and Eps15-DUB-dead (66.6%, 22.9s). n = 6 pits were analyzed for each plot. Data were shown as mean ± standard error. Scale bar = 1 μm. All cell FRAP experiments were performed at 37°C. **h**, Schematic showing how polyubiquitin stabilizes the endocytic protein network by interacting with and cross-linking UIMs on endocytic proteins, resulting in productive clathrin-mediated endocytosis (Top). Removal of ubiquitin from the endocytic protein network using DUB decreases the network multivalency thus making the network less stable, resulting in less efficient clathrin-mediated endocytosis (bottom).

### Light-activated recruitment of DUBs demonstrates that loss of ubiquitination destabilizes endocytic sites within minutes

While the results presented so far illustrate the importance of ubiquitylation to assembly of flexible protein networks at endocytic sites, it is not clear over what timescale these effects occur. In particular, does loss of ubiquitin acutely influence individual endocytic events or does it cause broader physiological changes that ultimately impact endocytosis? To address this question, we developed a system that allowed us to measure endocytic dynamics immediately after recruitment of deubiquitylases to endocytic sites. For this purpose, we made use of our previous observation that monomeric Eps15, which is created by deletion of Eps15’s coiled-coil domain, is not stably recruited to endocytic sites, likely owing to reduced affinity for the endocytic protein network (12). Therefore, we created a chimeric protein in which Eps15’s coiled-coil domain was replaced by a domain that forms dimers and oligomers upon blue light exposure, the photolyase homology region (PHR) of CRY2 (51) (Figure 6a). As reported previously, the resulting chimera, Eps15-CRY2 assembles upon blue light exposure, resulting in its stable recruitment to endocytic sites (12).

**Figure 6.**
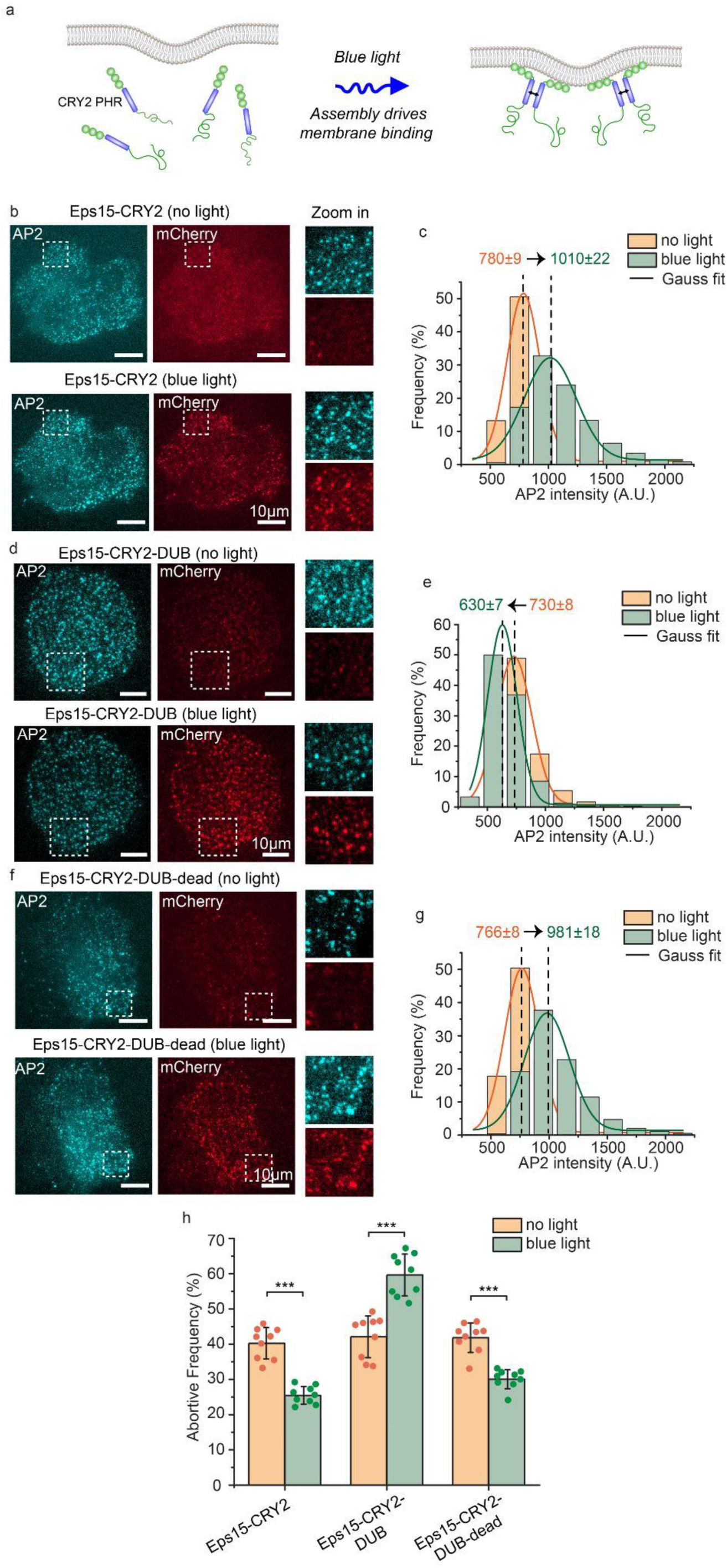
Light-activated recruitment of DUBs demonstrates that loss of ubiquitination destabilizes endocytic sites within minutes. **a**, Schematic of blue light driving assembly and membrane binding of Eps15-CRY2 chimera in which the Eps15 coiled-coil domain is replaced with the light-activation CRY2 PHR domain. **b, d, f**, Representative images of Eps15KO SUM cells expressing Eps15-CRY2 (**b**), Eps15-CRY2-DUB (**d**) and Eps15-CRY2-DUB-dead (**f**) before and after applying blue light. AP-2 σ2-HaloTag was labeled with JF646 (cyan). Insets show the zoom-in area of the white dashed box. mCherry was fused to all three constructs at their C terminus for visualization. Scale bar = 10 μm. **c, e, g**, Histograms and the Gaussian fit of the AP2 intensity distribution tracked in endocytic pits when expressing Eps15-CRY2 (**c**), Eps15-CRY2-DUB (**e**) and Eps15-CRY2-DUB-dead (**g**) before and after exposed to blue light, respectively. **h**, Frequency of short-lived structures comparison before and after blue light was applied to the cells under each condition. For Eps15-CYR2, n = 9 biologically independent cell samples were collected and in total 7539 pits (before light) and 7616 pits (blue light) were analyzed. For Eps15-CRY2-DUB, n = 9 and 7533 pits (before light) and 6626 pits (blue light) were analyzed. For Eps15-CRY2-DUB-dead, n = 9 and total pits = 8114 (before light) and 8514 (blue light). Dots represent frequency from each sample. An unpaired, two-tailed student’s t test was used for statistical significance. ***: P < 0.001. Error bars represent standard deviation. Cells were imaged at 37°C for all conditions.

Similarly, we found that when Eps15-CRY2 was transiently expressed in Eps15 knockout cells, it colocalized weakly with endocytic sites prior to blue light exposure (Figure 6b, top). The fraction of short-lived endocytic sites in these cells remained similar to the level in knockout cells (40.2 ± 4.4%, Figure 6h), consistent with previous findings that a monomeric version of Eps15 cannot rescue Eps15 knockout (12). However, upon exposure to blue light, Eps15-CRY2 was recruited to endocytic sites, and the fraction of pits showing Eps15 colocalization increased from 35.5 ± 5.5% to 74.1 ± 6%, Figure 6b, bottom; Figure S8). Simultaneously with the increase in Eps15 recruitment, more AP2 was recruited to endocytic sites (mean AP2 intensity shifted from 780 ± 9 to 1010 ± 22, Figure 6b, c, Supplementary Information Movie S3), suggesting increased stability and maturation of endocytic sites. Similarly, the fraction of short-lived endocytic structures was reduced to near wild type levels, 25.4 ± 2.5% (Figure 6h), confirming that light-induced assembly of Eps15 stabilized endocytic sites, rescuing the defects associated with Eps15 knockout (12). Importantly, in these experiments, endocytic dynamics were consecutively measured before and after blue light exposure in each cell, such that changes associated with light-activated protein assembly were directly observed for individual cells (see Materials and Methods).

We next repeated these experiments in Eps15 knockout cells that transiently expressed a blue light activated Eps15 chimera fused to the UL36 deubiquitylase enzyme, Eps15-CRY2-DUB. Similar to Eps15-CRY2, in the absence of blue light, this protein colocalized weakly with endocytic sites (Figure 6d, top, Figure S8) and failed to substantially reduce the fraction of short-lived endocytic sites (42.1 ± 5.9%, Figure 6h). This result confirms that the DUB enzyme has minimal impact on endocytosis prior to its recruitment to endocytic sites. However, upon exposure to blue light, increased recruitment of this DUB-containing chimera to endocytic sites resulted in a nearly 1.5-fold increase in the fraction of short-lived endocytic sites (59.6 ± 5.9% vs. 42.1 ± 5.9%, Figure 6h), which correlated with a substantial reduction in recruitment of AP2 to endocytic sites (730 ± 8 to 630 ± 7, Figure 6e, Supplementary Information Movie S3). In contrast, expression of an Eps15-CRY2 chimera that contained the catalytically inactive DUB, Eps15-CRY2-DUB-dead, reduced the fraction of short-lived endocytic structures to near wild type levels (30.0 ± 2.7% vs. 25.4 ± 2.5%, Figure 6h) upon blue light exposure. Similarly, recruitment of AP2 to endocytic sites in cells expressing Eps15-CRY2-DUB-dead was similar to that in cells expressing Eps15-CRY2 (Figure 6f, Figure 6g, 766 ± 8 to 981 ± 18, and Supplementary Information Movie S3), suggesting that the DUB fusion did not sterically inhibit endocytic dynamics. Importantly, the fraction of short-lived endocytic events in cells expressing each of the CRY2 chimeras was similar to that in Eps15 knockout cells, suggesting that expression of Eps15-CRY2-DUB did not significantly impact endocytic dynamics prior to its light-activated recruitment to endocytic sites. Taken together, these results suggest that loss of ubiquitylation acutely destabilizes endocytic sites within minutes.

## Discussion

Here we demonstrate that ubiquitylation initiates assembly of the flexible network of early initiator proteins responsible for clathrin-mediated endocytosis. Our *in vitro* experiments demonstrate that polyubiquitin stabilizes liquid-like droplets of Eps15, substantially raising the melting temperature of the resulting protein network. Importantly, these effects required Eps15’s ubiquitin interacting motif (UIM). Similarly, in live cell imaging experiments, a version of Eps15 lacking the UIM domain failed to rescue the increase in short-lived endocytic structures resulting from Eps15 knockout. These results suggest that interactions between Eps15 and ubiquitylated proteins, either transmembrane cargo or other endocytic proteins, stabilize early endocytic assemblies by promoting condensation of Eps15 proteins into liquid-like networks.

To test the impact of ubiquitin on the stability of endocytic sites more broadly, we evaluated the impact of deubiquitylase (DUB) enzymes on coated vesicle dynamics. Here we found that expressing a version of Eps15 fused to a broad-spectrum DUB substantially increased the number of unstable, short-lived endocytic sites, rather than rescuing the defect created by Eps15 knockout. Further, FRAP experiments, both on protein droplets *in vitro* and in live cells, demonstrated that interactions between Eps15 and ubiquitin decrease the molecular exchange of Eps15 within the protein network. Specifically, mutations that interfered with Eps15-ubiquitin interactions, such as deleting Eps15’s UIM domain or fusing a DUB domain to Eps15, increased exchange of Eps15. These data suggest that interactions with ubiquitin are necessary for stable recruitment of Eps15 to endocytic sites. In the context of a growing endocytic structure, interactions between ubiquitin and Eps15 may function to stabilize endocytic sites that contain ubiquitinated cargo, promoting robust, cargo-dependent endocytosis. Finally, by using a light activated recruitment system for Eps15-DUB, we demonstrated that loss of ubiquitin acutely destabilizes endocytic sites within minutes, a result which likely represents the cumulative effect of removing ubiquitin from multiple endocytic proteins and transmembrane protein cargos. Notably, while our *in vitro* experiments suggest that poly-ubiquitin has a more potent impact on Eps15 assembly than mono-ubiquitin, we speculate that a similar multi-valent effect could be achieved by mono-ubiquitination of multiple proteins that interact with Eps15, *in vivo*.

Our finding that polyubiquitin can stabilize early endocytic networks is supported by earlier work suggesting the importance of ubiquitylation during endocytosis (19, 52, 53). Specifically, Eps15 and Epsin are both known to contain UIM motifs (19, 54) and have long been thought to act as cargo adaptors for ubiquitinated transmembrane proteins (9, 54, 55). More recently, several laboratories reported that Eps15 and its homologs provide a flexible, liquid-like platform for endocytic assembly (12, 13, 15). These findings highlight the role of dynamic networks of intrinsically disordered proteins during endocytosis. Similarly, condensates have been found to play a role in membrane budding events throughout the cell including Golgi to plasma membrane transport (56), endophilin-mediated endocytosis (57, 58), and ER to Golgi transport (59). However, a key question has remained unanswered – how is the assembly of endocytic condensates regulated? Our work addresses this question by demonstrating that ubiquitylation, inherently a multi-valent process, stabilizes multi-valent interactions among early endocytic proteins, providing a previously unknown axis of dynamic control over endocytic initiation. Prior work on endocytic dynamics has suggested that nascent endocytic sites mature into productive endocytic structures by passing through a series of biochemical “checkpoints” or criteria, the precise identity of which remains unknown (1). In this context, our results suggest that the ubiquitin content of endocytic sites, which stabilizes the flexible network of early endocytic proteins, may constitute such a checkpoint.

## Materials and Methods

### Reagents

Tris-HCl (Tris hydrochloride), HEPES (4-(2-hydroxyethyl)-1-piperazineethanesulfonic acid), IPTG (isopropyl-β-D-thiogalactopyranoside), NaCl, β-mercaptoethanol, Triton X-100, neutravidin, and Texas Red-DHPE (Texas Red 1,2-dihexadecanoyl-sn-glycero-3-phosphoethanolamine triethylammonium salt) were purchased from Thermo Fisher Scientific. Sodium bicarbonate, sodium tetraborate, EDTA (Ethylene diamine tetraacetic acid), EGTA (Ethylene glycol tetraacetic acid), glycerol, TCEP (tris(2-carboxyethyl) phosphine), DTT (Dithiothreitol), PMSF (phenylmethanesulfonylfluoride), EDTA-free protease inhibitor tablets, thrombin, imidazole, sodium bicarbonate, PLL (poly-l-lysine), Atto640 NHS ester and Atto488 NHS ester were purchased from Sigma-Aldrich. Human Holo-Transferrin protein, Monoubiquitin and K63 linked Tetraubiquitin were purchased from Boston Biochem (Catalog #: 2914-HT-100MG, U-100H, and UC-310). PEG 8000 (Polyethylene glycol 8000) was purchased from Promega (Catalog #: V3011). Amine-reactive PEG (mPEG-succinimidyl valerate MW 5000) and PEG-biotin (Biotin-PEG SVA, MW 5000) were purchased from Laysan Bio. DP-EG10-biotin (dipalmitoyl-decaethylene glycol-biotin) was provided by D. Sasaki (Sandia National Laboratories). POPC (1-palmitoyl-2-oleoyl-glycero-3-phosphocholine) and DGS-NTA-Ni (1,2-dioleoyl-sn-glycero-3-[(N-(5-amino-1-carboxypentyl) iminodiacetic acid)-succinyl] (nickel salt)) were purchased from Avanti Polar Lipids. All reagents were used without further purification.

### Plasmids

Plasmids used for purifying Eps15 and Eps15ΔUIM from bacteria are pET28a-6×His-Eps15 (FL) and pET28a-6×His-Eps15ΔUIM. pET28a-6×His-Eps15 (FL) encoding H. sapiens Eps15 was kindly provided by T. Kirchhausen (26), Harvard Medical School, USA. pET28a-6×His-Eps15ΔUIM was generated by using site-directed mutagenesis to introduce a stop codon after residue 850 of Eps15 to generate a truncated version lacking residues 851-896 corresponding to the UIM domains at the C terminus. The forward primer 5’-GTGCTTATCCCTGAGAAGAAGATATGATCG-3’ and reverse primer 5’-CATATCTTCTTCTCAGGGATAAGCACTGAAG-3’ were used.

Plasmids used for mammalian cell expression of Eps15 variants were derived from Eps15-pmCherryN1 (Addgene plasmid #27696, a gift from C. Merrifield (60)), which encodes Eps15-mCherry (denoted as Eps15wt). All of the following Eps15 variants contain mCherry at their C terminal end for visualization even though mCherry is not mentioned in their names. The Eps15ΔUIM plasmid was generated by PCR-mediated deletion of the UIM domains (residues 851-896). The 138 base pairs corresponding to the two UIM domains were deleted using the 5’ phosphorylated forward primer 5’-CGGATGGGTTCGACCTC-3’ and the reverse primer 5’-GGGATAAGCACTGAAGTTGG-3’. After PCR amplification and purification, the PCR product was recircularized to generate the Eps15ΔUIM. Plasmids encoding the broad-spectrum deubiquitylase (DUB), UL36, and the catalytically inert mutant (C56S, DUB-dead), were generously provided by J. A. MacGurn (42). Plasmids encoding the Eps15-DUB and Eps15-DUB-dead were generated by restriction cloning. Amplification of the DUB and DUB-dead was achieved using the forward primer 5’-CATGAGGATCCAATGGACTACAAAGACCATGACG-3’ and the reverse primer 5’-CATGAGGATCCGGGTATGGGTAAAAGATGCGG-3’. The amplicon was then inserted into the Eps15-pmCherryN1 at the BamH1 restriction sites between Eps15 and mCherry. The plasmid encoding Eps15-CRY2 was generated by using the crytochrome 2 photolyase homology region (CRY2 PHR) domain of *Arabidopsis thaliana* to replace the coiled-coil domain in Eps15 based on our previous report (12). Specifically, the CRY2 PHR domain was PCR amplified from pCRY2PHR-mCherryN1 (Addgene plasmid #26866, a gift from C. Tucker (61)) using primers 5′-TAGGATCAAGTCCTGTTGCAGCCACCATGAAGATGGACAAAAAGAC-3′ and 5′-ATCAGTTTCATTTGCATTGAGGCTGCTGCTCCGATCAT-3′. This fragment was inserted by Gibson assembly (New England Biolabs) into Eps15-pmCherryN1 (Addgene plasmid#27696, a gift from C. Merrifield), which were PCR amplified to exclude Eps15 coiled-coil domain (residues 328–490) using primers 5′-TCATGATCGGAGCAGCAGCCTCAATGCAAATGAAACTGATGGAAATGAAAGATTTGGAAAATCATAATAG-3′ and 5′-TTGTCCATCTTCATGGTGGCTGCAACAGGACTTGATCCTATGAT-3′. Plasmids encoding Eps15-CRY2-DUB and Eps15-CRY2-DUB-dead were generated by inserting DUB and DUB-dead in between Eps15-CRY2 and mCherry through Gibson assembly. The primers 5’-GTCAGCTGGCCCGGGATCCAATGGACTACAAA-3’ and 5’-CTCACCATGGTGGCGACCGGTGGATCCGGGTA-3’ were used for amplifying DUB and DUB-dead and primers 5’-TTTACCCATACCCGGATCCACCGGTCGCCACCA-3’ and 5’-TCATGGTCTTTGTAGTCCATTGGATCCCGGGCCAG-3’ were used for amplifying the vector Eps15-CRY2.

All constructs were confirmed by DNA sequencing.

### Protein purification

Eps15 and Eps15ΔUIM were purified based on a previously reported protocol (12). Briefly, full-length Eps15 and Eps15ΔUIM were expressed as N-terminal 6×His-tagged constructs in BL21 (DE3) Escherichia coli cells. Cells were grown in 2×YT medium for 3-4 h at 30 °C to an optical density at 600 nm of 0.6-0.9, then protein expression was induced with 1 mM IPTG at 30 °C for 6-8 hours. Cells were collected, and bacteria were lysed in lysis buffer using homogenization and probe sonication on ice. Lysis buffer consisted of 20 mM Tris-HCl, pH 8.0, 300 mM NaCl, 5% glycerol, 10 mM imidazole, 1 mM TCEP, 1 mM PMSF, 0.5% Triton X-100 and 1 EDTA-free protease inhibitor cocktail tablet (Roche: Cat#05056489001) per 50 mL buffer. Proteins were incubated with Ni-NTA Agarose (Qiagen, Cat#30230) resin for 30 min at 4 °C in a beaker with stirring, followed by extensive washing with 10 column volumes of lysis buffer with 20 mM imidazole and 0.2% Triton X-100 and 5 column volumes of buffer without Triton X-100. Then proteins were eluted from the resin in 20 mM Tris-HCl, pH 8.0, 300 mM NaCl, 5% glycerol, 250 mM imidazole, 0.5 mM TCEP, 1 mM PMSF, and EDTA-free protease inhibitor cocktail tablet. Eluted proteins were further purified by gel filtration chromatography using a Superose 6 column run in 20 mM Tris-HCl, pH 8.0, 150 mM NaCl, 1 mM EDTA, and 5 mM DTT. For droplet experiments, prior to running the gel filtration column, the 6×His tag on the proteins was further cleaved with Thrombin CleanCleave kit (Sigma-Aldrich, Cat# RECMT) overnight at 4 °C on the rocking table after desaltig in 50 mM Tris-HCl, pH 8.0, 10 mM CaCl_2_, 150 mM NaCl and 1 mM EDTA using a Zeba Spin desalting column (Thermo Scientific, Cat#89894). The purified proteins were dispensed into small aliquots, flash frozen in liquid ntitrogen and stored at –80 °C.

### Protein labeling

Eps15 and Eps15ΔUIM were labeled with amine-reactive NHS ester dyes Atto488 in phosphate-buffered saline (PBS, Hyclone) containing 10 mM sodium bicarbonate, pH 8.3. Monoubiquitin and tetraubiquitin were labeled with Atto640 in PBS, pH 7.4. The concentration of dye was adjusted experimentally to obtain a labeling ratio of 0.5–1 dye molecule per protein, typically using 2-fold molar excess of dye. Reactions were performed for 30 min on ice. Then labeled Eps15 and Eps15ΔUIM was buffer exchanged into 20 mM Tris-HCl pH 7.5, 150 mM NaCl, 5 mM TCEP, 1 mM EDTA, 1 mM EGTA and separated from unconjugated dye using Princeton CentriSpin-20 size-exclusion spin columns (Princeton Separations). The labeled monoubiquitin and tetraubiquitin were buffer exchanged to 20 mM Tris-HCl, 150 mM NaCl, pH 7.5 and separated from unconjugated dye as well using 3K Amicon columns. Labeled proteins were dispensed into small aliquots, flash frozen in liquid nitrogen and stored at –80°C. For all experiments involving labeled Eps15 and Eps15ΔUIM, a mix of 90% unlabelled/10% labeled protein was used. 100% labeled monoubiquitin and tetraubiquitin were directly used due to their small fraction compared to Eps15 variants.

### PLL-PEG preparation

PLL-PEG and biotinylated PLL-PEG were prepared as described previously with minor alterations (62). Briefly, for PLL-PEG, amine-reactive mPEG-succinimidyl valerate was mixed with poly-L-lysine (15–30 kD) at a molar ratio of 1:5 PEG to poly-L-lysine. For biotinylated PLL-PEG, amine reactive PEG and PEG-biotin was first mixed at a molar ratio of 98% to 2%, respectively, and then mixed with PLL at 1:5 PEG to PLL molar ratio. The conjugation reaction was performed in 50 mM sodium tetraborate pH 8.5 solution and allowed to react overnight at room temperature with continued stirring. The products were buffer exchanged into 5 mM HEPES, 150 mM NaCl pH 7.4 using Zeba spin desalting columns (7K MWCO, ThermoFisher) and stored at 4 °C.

### Protein droplets

Eps15 or Eps15ΔUIM droplets were formed by mixing proteins with 3% w/v PEG8000 in 20 mM Tris-HCl pH 7.5, 150 mM NaCl, 5 mM TCEP 1 mM EDTA, 1 mM EGTA. 7 μM Eps15 or Eps15ΔUIM was used to form the droplets, with addition of ubiquitins accordingly. For imaging droplets, 2% PLL-PEG were used to passivate coverslips (incubated for 20 min) before adding protein-containing solutions. Imaging wells consisted of 5 mm diameter holes in 0.8 mm thick silicone gaskets (Grace Bio-Labs). Gaskets were placed directly onto no.1.5 glass coverslips (VWR International), creating a temporary water-proof seal. Prior to well assembly, gaskets and cover slips were cleaned in 2% v/v Hellmanex III (Hellma Analytics) solution, rinsed thoroughly with water, and dried under a nitrogen stream. The imaging well was washed 6-8 times with 20 mM Tris-HCl, 150 mM NaCl and 5 mM TCEP buffer before adding solutions that contained proteins.

### GUV preparation

GUVs consisted of 93 mol% POPC, 5 mol% Ni-NTA, and 2 mol% DP-EG10 biotin. GUVs were prepared by electroformation according to published protocols (63). Briefly, lipid mixtures dissolved in chloroform were spread into a film on indium-tin-oxide (ITO) coated glass slides (resistance ∼8-12 W per square) and further dried in a vacuum desiccator for at least 2 hours to remove all of the solvent. Electroformation was performed at 55°C in glucose solution with an osmolarity that matched the buffer to be used in the experiments. The voltage was increased every 3 min from 50 to 1400 mV peak to peak for the first 30 min at a frequency of 10 Hz. The voltage was then held at 1400 mV peak to peak, 10 Hz, for 120 min and finally was increased to 2200 mV peak to peak, for the last 30 min during which the frequency was adjusted to 5 Hz. GUVs were stored in 4°C and used within 3 days after electroformation.

### GUV tethering

GUVs were tethered to glass coverslips for imaging as previously described (64). Briefly, glass cover slips were passivated with a layer of biotinylated PLL-PEG, using 5 kDa PEG chains. GUVs doped with 2 mol% DP-EG10-biotin were then tethered to the passivated surface using neutravidin. Imaging wells consisted of 5 mm diameter holes in 0.8 mm thick silicone gaskets were prepared by placing silicone gaskets onto Hellmanex III cleaned coverslips. In each imaging well, 20 μL of biotinylated PLL-PEG was added. After 20 min of incubation, wells were serially rinsed with appropriate buffer by gently pipetting until a 15,000-fold dilution was achieved. Next, 4 μg of neutravidin dissolved in 25 mM HEPES, 150 mM NaCl (pH 7.4) was added to each sample well and allowed to incubate for 10 minutes. Wells were then rinsed with the appropriate buffer to remove excess neutravidin. GUVs were diluted in 20 mM Tris-HCl, 150 mM NaCl, 5 mM TCEP, pH 7.5 at ratio of 1:13 and then 20 μL of diluted GUVs was added to the well and allowed to incubate for 10 minutes. Excess GUVs were then rinsed from the well using the same buffer and the sample was subsequently imaged using confocal fluorescence microscopy.

### Cell culture

Human-derived SUM159 cells gene-edited to add a HaloTag to both alleles of AP-2 σ2 were a gift from T. Kirchhausen (36). Cells were further gene-edited to knock out both alleles of endogenous Eps15 using CRISPR-associated protein 9 (Cas9) to produce the Eps15 knockout cells developed previously by our group (12).

Cells were grown in 1:1 DMEM high glucose: Ham’s F-12 (Hyclone, GE Healthcare) supplemented with 5% fetal bovine serum (Hyclone), Penicillin/Streptomycin/l-glutamine (Hyclone), 1 μg ml^−1^ hydrocortisone (H4001; Sigma-Aldrich), 5 μg ml^−1^ insulin (I6634; Sigma-Aldrich) and 10 mM HEPES, pH 7.4 and incubated at 37 °C with 5% CO_2_. Cells were seeded onto acid-washed coverslips at a density of 3 × 10^4^ cells per coverslip for 24 h before transfection with 1μg of plasmid DNA using 3 μl Fugene HD transfection reagent (Promega). HaloTagged AP-2 σ2 was visualized by adding Janelia Fluor 646-HaloTag ligand (Promega). Ligand (100 nM) was added to cells and incubated at 37 °C for 15 min. Cells were washed with fresh medium and imaged immediately.

### Fluorescence microscopy

Images of protein droplets and GUVs were collected on a spinning disc confocal super resolution microscope (SpinSR10, Olympus, USA) equipped with a 1.49 NA/100X oil immersion objective. For GUV imaging, image stacks taken at fixed distances perpendicular to the membrane plane (0.5 μm steps) were acquired immediately after GUV tethering and again after protein addition. Images taken under deconvolution mode were processed by the built-in deconvolution function in Olympus CellSens software (Dimension 3.2, Build 23706). At least 30 fields of views were randomly selected for each sample for further analysis. Imaging was performed 5 min after adding proteins, providing sufficient time to achieve protein binding and reach a steady state level of binding. For experiments used to construct phase diagrams, temperature was monitored by a thermistor placed in the protein solution in a sealed chamber to prevent evaporation. Samples were heated from room temperature through an aluminum plate fixed to the top of the chamber. Temperature was increased in steps of 1 °C until the critical temperature was reached. Images of at least 20 droplets were taken at each temperature, once the temperature stabilized.

Live-cell images were collected on a TIRF microscope consisting of an Olympus IX73 microscope body, a Photometrics Evolve Delta EMCCD camera, and an Olympus 1.4 NA ×100 Plan-Apo oil objective, using MicroManager version 1.4.23. The coverslip was heated to produce a sample temperature of 37 °C using an aluminum plate fixed to the back of the sample. All live-cell imaging was conducted in TIRF mode at the plasma membrane 48 h after transfection. Transfection media used for imaging lacked pH indicator (phenol red) and was supplemented with 1 μL OxyFluor (Oxyrase, Mansfield, OH) per 33 μL media to decrease photobleaching during live-cell fluorescence imaging. 532 nm and 640 nm lasers were used for excitation of mCherry and Janelia Fluor 646-HaloTag ligand of AP2, respectively. Cell movies were collected over 10 min at 2 s intervals between frames. For blue-light assays, samples were exposed to 25 μW 473 nm light as measured out of the objective when in wide-field mode. Blue light was applied for 500 ms every 3 s for cell samples. Cell movies were collected over 11 min at 3 s intervals between frames, and analysis of movies began after 1 min of imaging to allow for blue light to take effect.

Droplet and live-cell FRAP experiments were performed using Olympus FRAP unit 405 nm laser on the spinning disc confocal microscope. Images were acquired every 3 s for 3 min after photobleaching. All droplet FRAP experiments were conducted at room temperature while live cell FRAP experiments were done at 37 °C.

### Image analysis

Fluorescence images analyzed in ImageJ (http://rsbweb.nih.gov/ij/). Intensity values along line scans were measured in unprocessed images using ImageJ. For phase diagrams, fluorescence intensity was measured in the center square of a 3 × 3 grid for each image where illumination was even. Two images were analyzed at each temperature for each condition. The intensity was normalized to the mean intensity value of the solution (200 A.U.).

Droplet FRAP data were analyzed using the FRAP Profiler plugin (https://worms.zoology.wisc.edu/research/4d/4d.html) for ImageJ. Droplets of similar size with similarly sized photobleached regions iwere selected for FRAP analysis. A single exponential fit was selected in the FRAP Profiler to fit the data and derive the mobile fraction and t_1/2_ for each droplet. Mean and standard deviation of mobile fraction and t_1/2_ were calculated for each condition.

Cell FRAP data were manually analyzed due to the high mobility of the endocytic structures on the plasma membrane. The intensity range was normalized to maximum and minimum intensity values, corresponding to pre– and post-bleaching frames. The mobile fraction and t_1/2_ were derived from the single exponential fit of the fluorescence recovery plot for each condition.

Clathrin-coated structures were detected and tracked using cmeAnalysis in MATLAB (65). The point spread function of the data was used to determine the standard deviation of the Gaussian function. AP-2 σ2 signal was used as the master channel to track clathrin-coated structures. Detected structures were analyzed if they persisted in at least three consecutive frames. The lifetimes of clathrin-coated that met these criteria were recorded for lifetime distribution analysis under different conditions.

### Flow cytometry

The internalization of transferrin through clathrin-mediated endocytosis was measured using flow cytometry based on previous reports (66–68). First, human holo-transferrin was labeled with atto-488 in PBS, pH 7.4 using the protein labeling methods above. Eps15 knockout SUM159 cells were plated on a 6-well plate at a density of 100,000 cells per well and a total volume of 2 mL per well. SUM159 cells endogenously expressing Eps15 were plated as a control. After 24 hr, media was aspirated from the cells and they were rinsed with PBS 3 times. The cells were starved for 30 min in 1mL pre-warmed serum-free media (SFM) and were then incubated with 50 μg/mL transferrin atto-488 in SFM for 30 min. Subsequently, the cells were incubated with 500 μL trypsin for about 1 min at 37°C, and then the trypsin was removed and replaced with fresh trypsin. This process was repeated 3x to wash away any surface bound transferrin. After the third 1 min trypsin wash, cells were fully detached from the wells by incubation with 500 μL of trypsin for 5 min at 37°C, 5% CO^2^. The trypsin was then quenched with 1 mL of media, and cells from each well were transferred to Eppendorf tubes and centrifuged at 300 x g for 5 min. The resulting cell pellet was resuspended in 300 μL of PBS in preparation for flow cytometry analysis.

A Guava easyCyte Flow Cytometer (Millipore Sigma) with 488 nm and 532 nm excitation lasers was used to analyze fluorescence of cells after transferrin incubation. All data were collected at 35 μL/min and flow cytometry data were analyzed using FlowJo (Treestar). A gate was drawn in forward scattering versus side scattering plots to exclude debris and contain the majority of cells (Figure S6). Within this gate, cell populations were further analyzed by plotting histograms of FITC fluorescence intensity to determine shifts in transferrin atto-488 uptake within cells from each group (Figure S7). Average fluorescence intensity was used to indicate the level of transferrin internalization.

### Statistical analysis

For all experiments yielding micrographs, each experiment was repeated independently on different days at least three times, with similar results. Phase diagram experiments were repeated independently twice with similar results. Collection of cell image data for clathrin-mediated endocytosis analysis was performed independently on at least two different days for each cell type or experimental condition. Statistical analysis was carried out using a two-tailed student’s t-test (unpaired, unequal variance) to probe for statistical significance (P < 0.05).

### Supplementary Information

**Movie S1: Eps15 droplets fusion with addition of MonoUb.** Eps15 (7μM) droplets (green) were incubated with 1μM MonoUb (magenta) in 20 mM Tris-HCl, 150 mM NaCl, 5 mM TCEP, 1 mM EDTA and 1 mM EGTA at pH 7.5 with 3% w/v PEG8000.

**Movie S2: Eps15 droplets fusion with addition of TetraUb.** Eps15 (7μM) droplets (green) were incubated with 0.25μM K63 linkage TetraUb (magenta) in 20 mM Tris-HCl, 150 mM NaCl, 5 mM TCEP, 1 mM EDTA and 1 mM EGTA at pH 7.5 with 3% w/v PEG8000.

**Figure S1.**
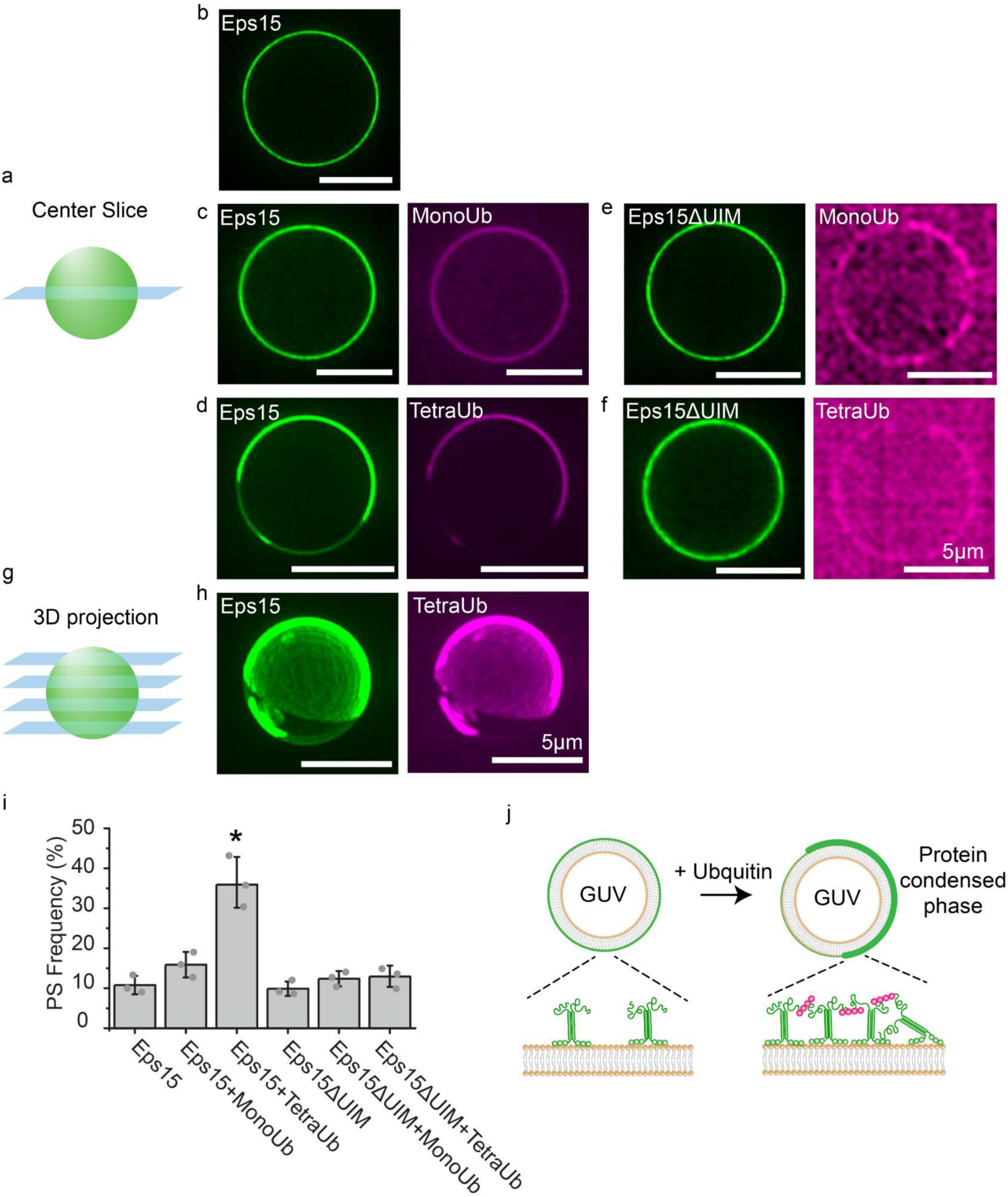
Polyubiquitin promotes phase separation of Eps15 on membrane surfaces. a, Cartoon depicting the center slice of a GUV and b-f, Representative center slice images of GUVs incubated with indicated proteins: 0.5 μM his-Eps15 alone (b), 0.5 μM his-Eps15 with 0.5 μM MonoUb (c), 0.5 μM his-Eps15 with 0.125 μM TetraUb (d), 0.5 μM his-Eps15ΔUIM with 0.5 μM MonoUb (e), and 0.5 μM his-Eps15ΔUIM with 0.125 μM TetraUb (f). g, Cartoon demonstrating the 3D reconstruction of a GUV, and h, the corresponding z-projections of the GUV from d, showing Eps15 assembled into protein condensed region together with TetraUb on GUV membrane. All scale bars are 5 μm. i, Frequency of GUVs displaying protein-rich domains for each set of proteins. GUVs were counted as displaying protein-rich domains if they contained distinct regions in which protein signal intensity differed by at least two-fold and the bright region covered at least 10% of the GUV surface in any z-slice. For each bar, n = 3 biologically independent experiments (each individual dot) with at least 44 total GUVs for each condition. Data are mean ± SD. *: P < 0.001 compared to all other groups using unpaired, two-tailed student’s t test. GUVs contain 93 mol% POPC, 5 mol% DGS-NTA-Ni, and 2 mol% DP-EG10-biotin. All experiments were conducted in 20 mM Tris-HCl, 150 mM NaCl, 5 mM TCEP at pH 7.5 buffer. j, Cartoon illustrating his-Eps15 binding to GUV membrane (left) and polyubiquitin driving Eps15 phase separation on GUV membrane by linking Eps15 through interaction with the UIMs (right).

**Figure S2.**
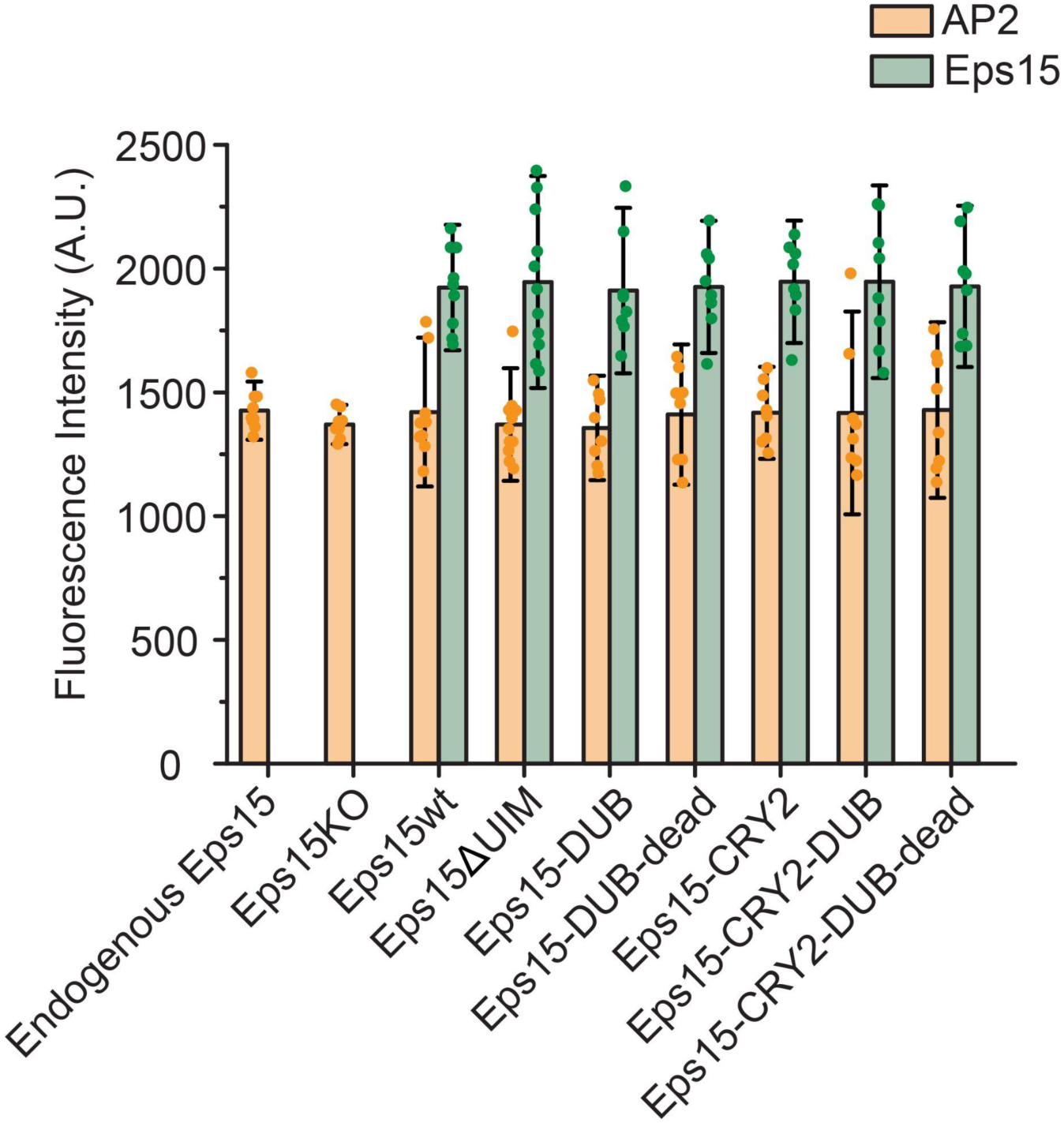
Eps15 mutants expression levels in each group measured from the mean fluorescence intensity of the cell images taken from TIRF microscopy. N = 9 biologically independent cells (each individual dot) was analyzed for each condition. Data are mean ± SD.

**Figure S3.**
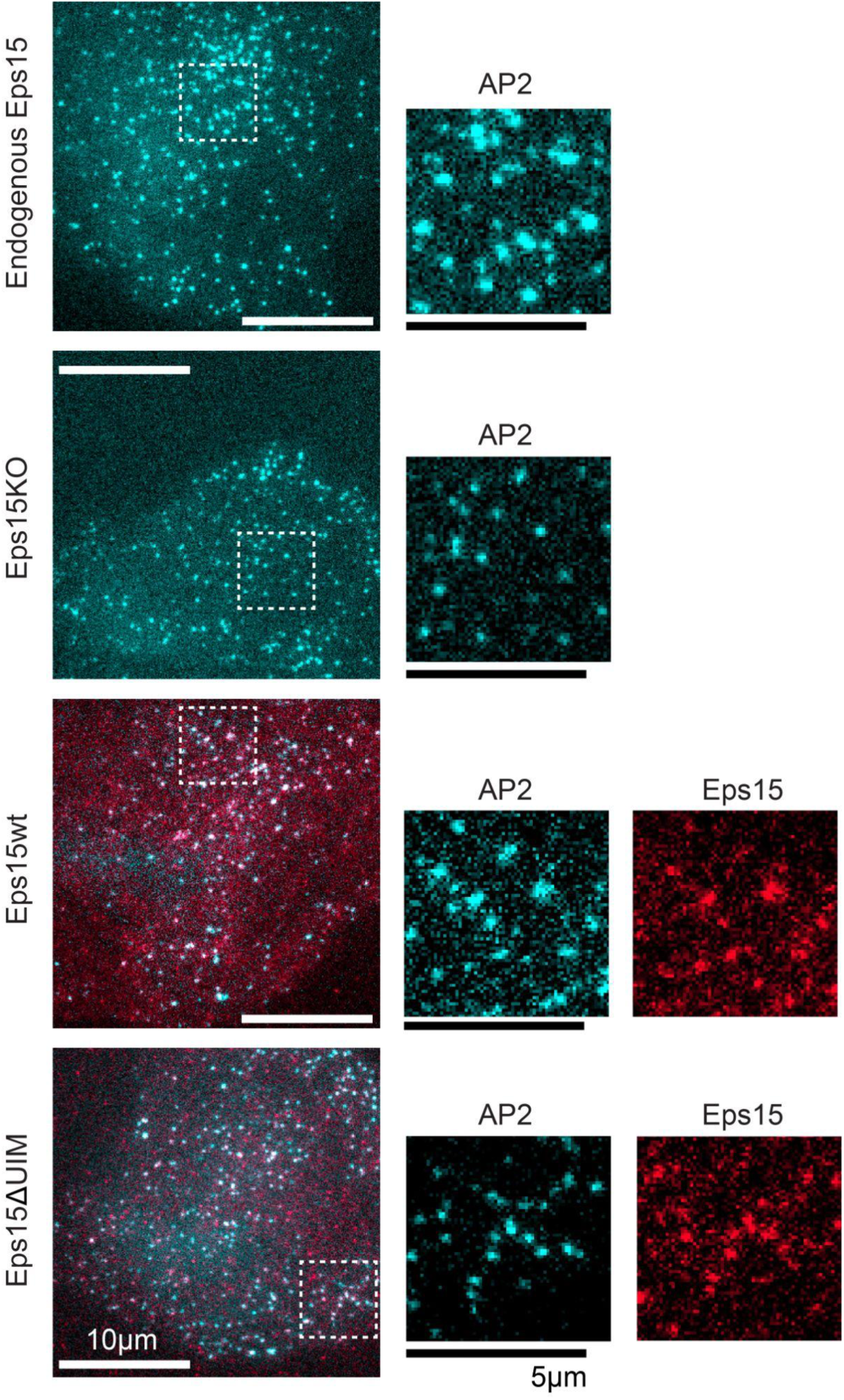
Representative image of endocytic pits in live SUM cells expressing gene-edited AP2 σ2-HaloTag: JF646 (cyan) and Eps15 variants (red). Endogenous Eps15 represents SUM cells expressing gene-edited AP2 σ2-HaloTag. Eps15KO represents SUM cells that were further CRISPR modified to knockout alleles of endogenous Eps15. Eps15wt and Eps15ΔUIM represent Eps15KO cells transfected with wildtype Eps15 and Eps15 with the depletion of both UIM domains, respectively. mCherry was fused to the C terminus of Eps15 and Eps15ΔUIM for visualization. Insets are the zoom-in area of the white dashed box. Scale bars are labeled in images. Cells were imaged at 37°C for all conditions.

**Figure S4.**
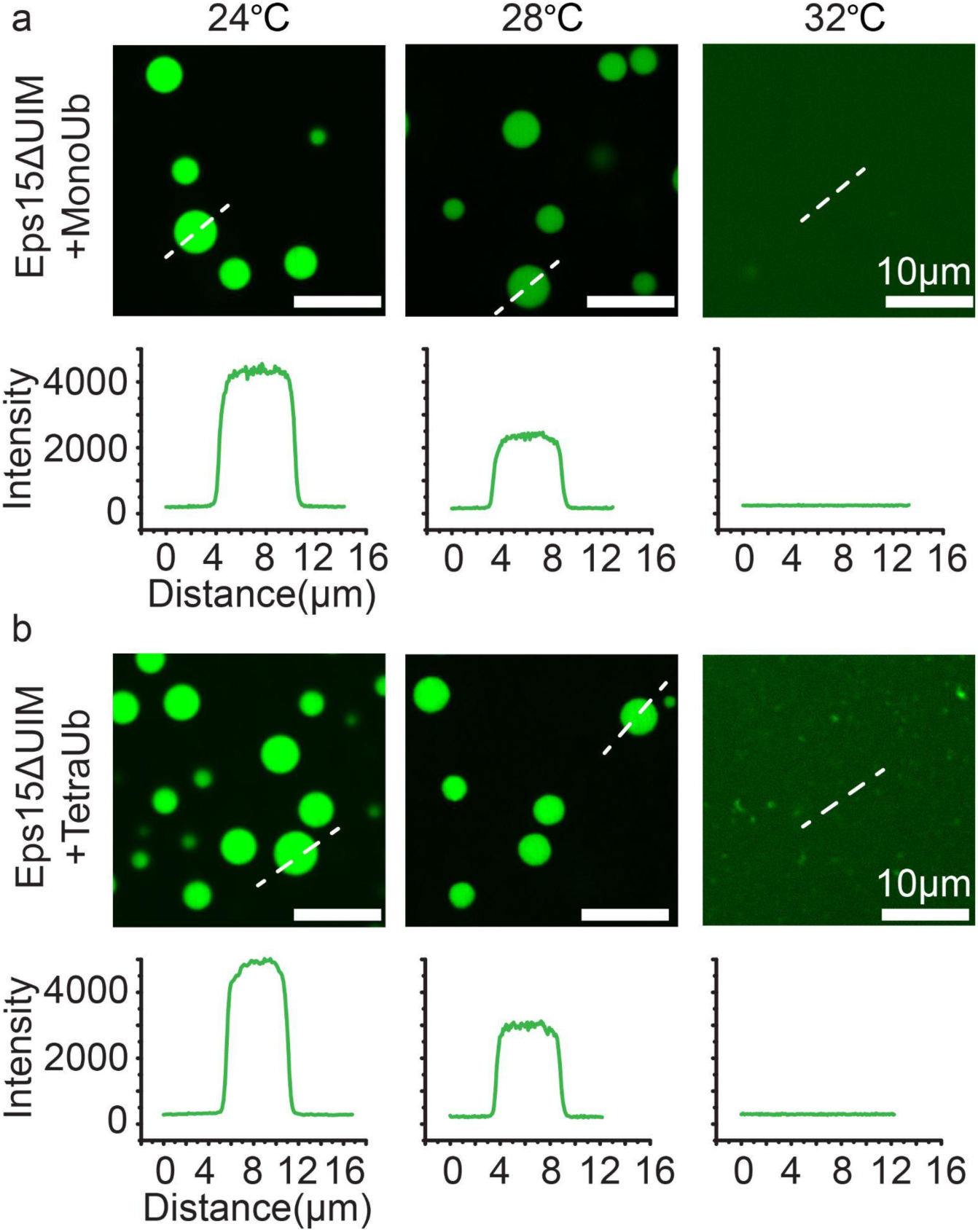
Representative images of protein droplets at increasing temperatures. Plots show fluorescence intensity of Eps15ΔUIM measured along dotted lines in each image. Droplets are formed from (**a**) 0.5 μM MonoUb, 7 μM Eps15ΔUIM and (**b**) 0.1 μM TetraUb, 7 μM Eps15ΔUIM in 20 mM Tris-HCl, 150 mM NaCl, 5 mM TCEP, 1 mM EDTA and 1 mM EGTA at pH 7.5 buffer with 3% PEG8000. Scale bars equal 10 μm.

**Figure S5.**
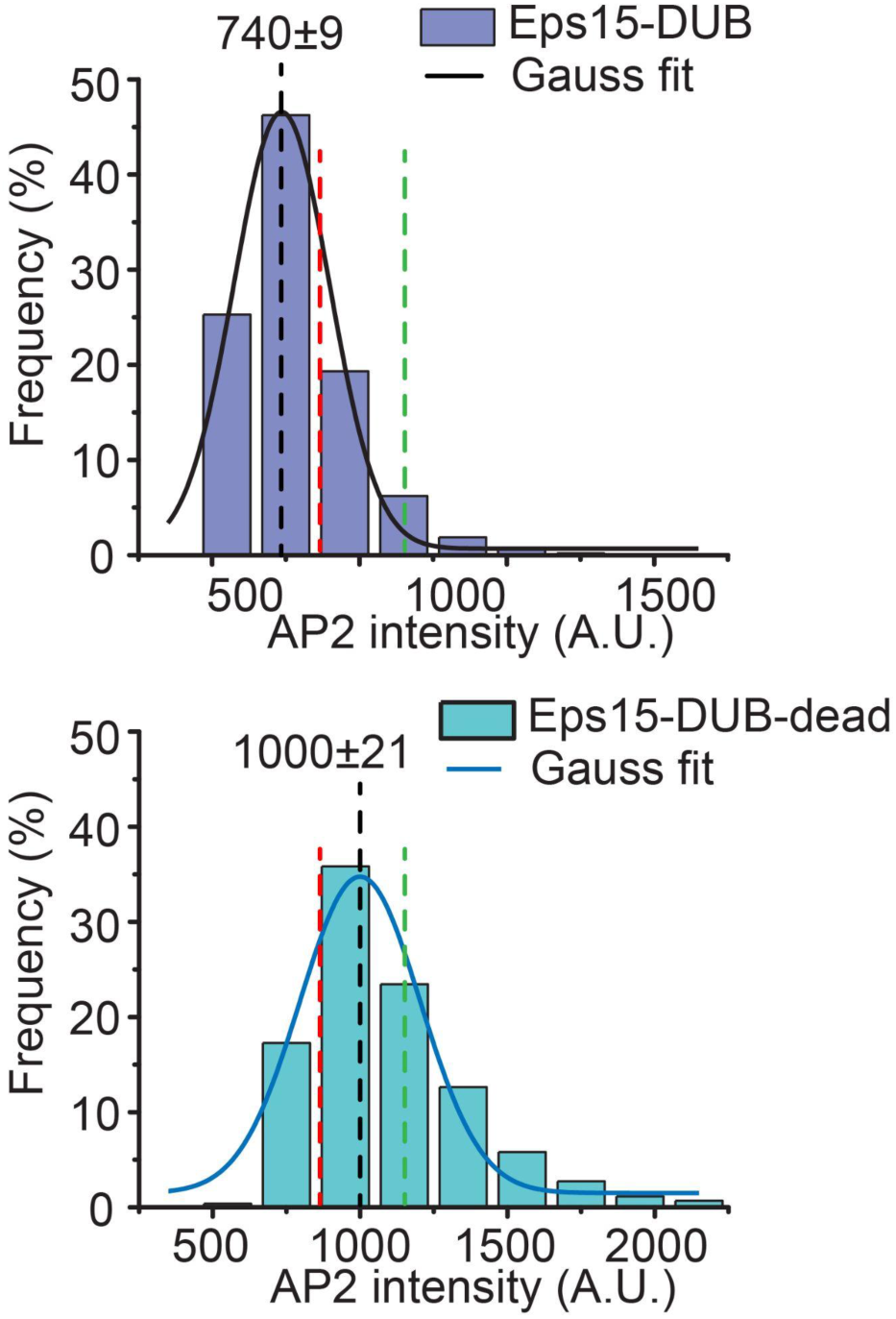
Histograms and the Guass fit of the AP2 intensity distribution tracked in endocytic pits when Eps15KO SUM cells were transfected to express Eps15-DUB (top) and Eps15-DUB-dead (bottom), respectively. The green dotted line and red dotted line are corresponding to endogenous Eps15 group and Eps15KO group in Figure 2. For Eps15-DUB, n = 9 biologically independent cell samples were collected and in total 8640 pits were analyzed. For Eps15-DUB-dead, n = 9 and 7420 pits were analyzed. Cells were imaged at 37°C.

**Figure S6.**
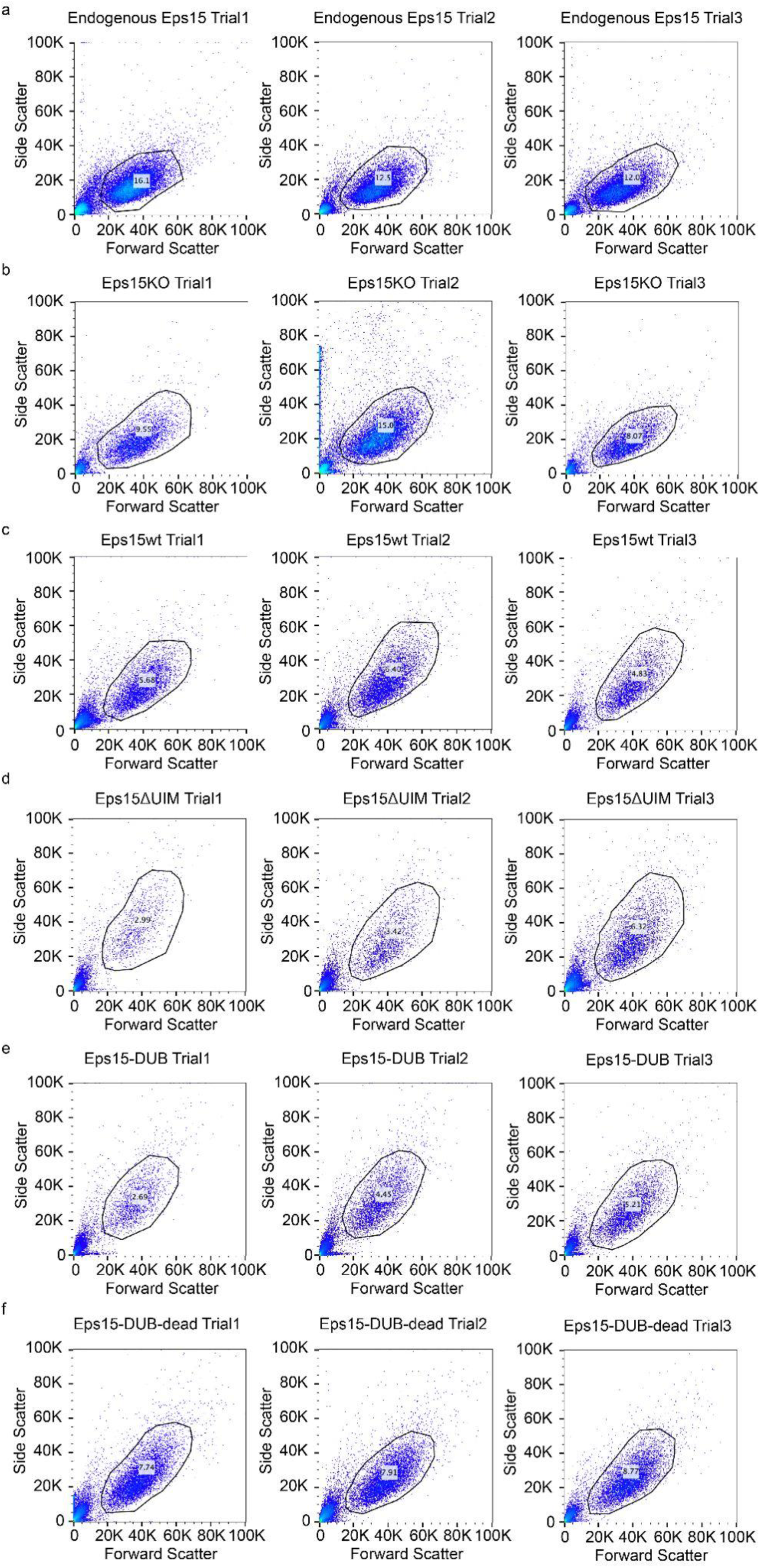
Flow cytometry scatterplots from populations of cells in each condition. The black irregular circle represents the gate. N=3 trials were did for each condition.

**Figure S7.**
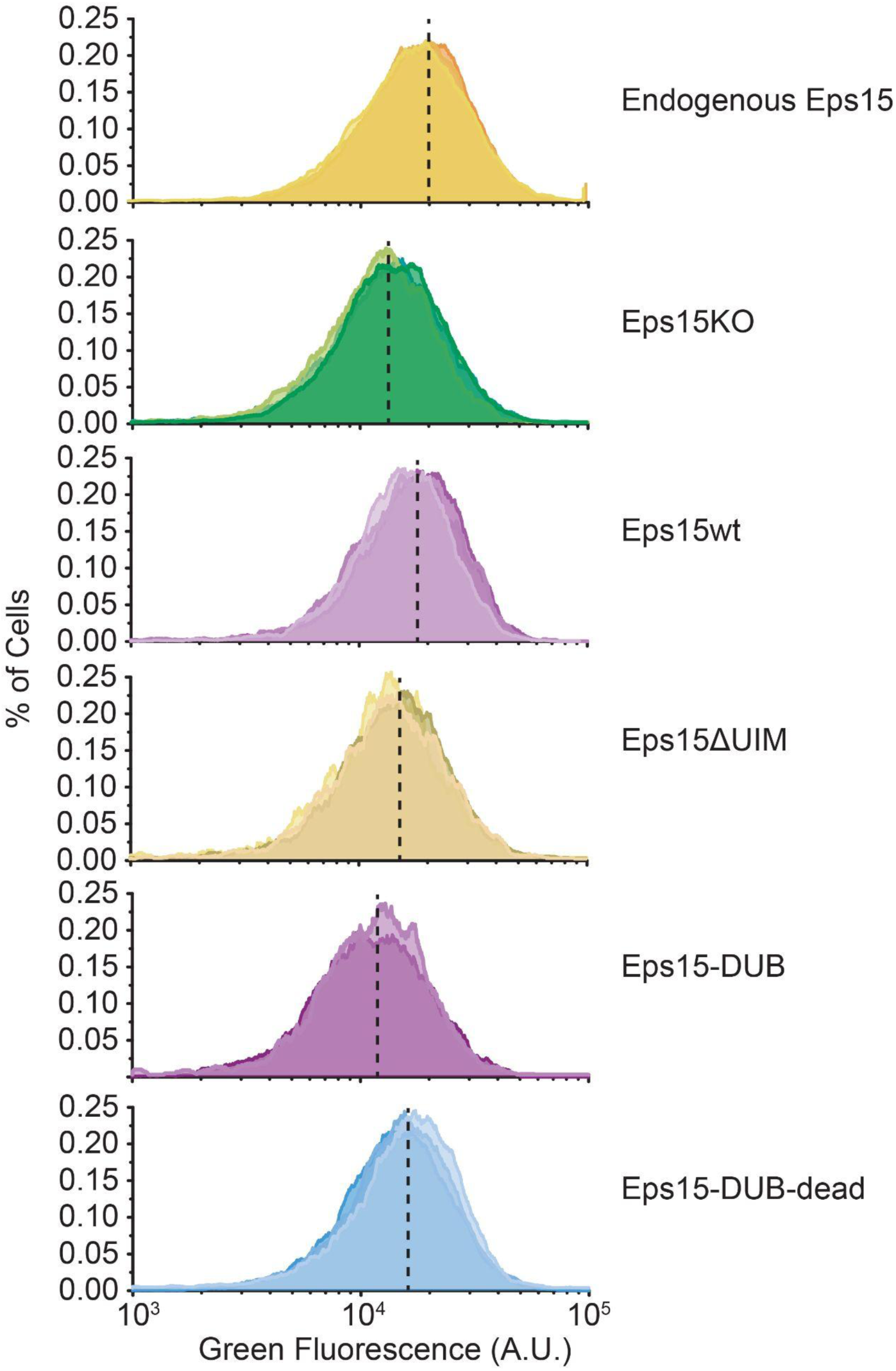
Flow cytometry histogram of the transferrin fluorescence intensity of cells in each condition. N=3 trials were did for each condition.

**Figure S8.**
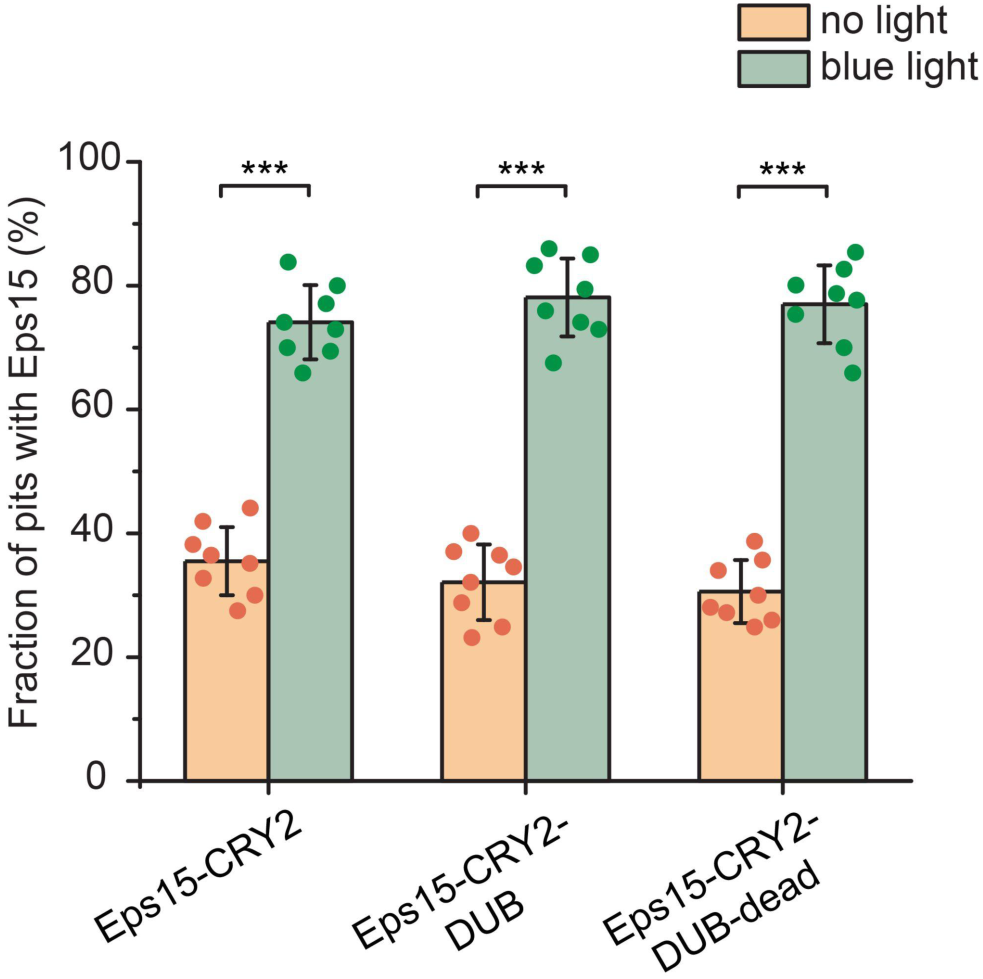
Fraction of endocytic pits showing Eps15 colocalization before and after exposure to blue light when expressing Eps15-CRY2, Eps15-CRY2-DUB, or Eps15-CRY2-DUB-dead. For Eps15-CYR2, n = 9 biologically independent cell samples were collected and in total 7539 pits (before light) and 7616 pits (blue light) were analyzed. For Eps15-CRY2-DUB, n = 9 and 7533 pits (before light) and 6626 pits (blue light) were analyzed. For Eps15-CRY2-DUB-dead, n = 9 and total pits = 8114 (before light) and 8514 (blue light). Dots represent frequency from each sample. An unpaired, two-tailed student’s t test was used for statistical significance. ***: P < 0.001. Error bars represent standard deviation. Cells were imaged at 37°C for all conditions.

**Movie S3**: Change of Eps15 and AP2 channel upon light activation in cells expressing Eps15-CRY2, Eps15-CRY2-DUB and Eps15-CRY2-DUB-dead, respectively.

## Acknowledgements

We thank T. Kirchhausen for the gift of SUM159/AP2-σ2-Halo Tag cells and J. MacGurn for the gift of plasmids encoding deubiquitylase UL36 and its catalytically inactive mutant. This research was supported by the National Institutes of Health through grants R35GM139531 to J.C.S., the National Science Foundation Directorate of Biological Sciences through grant 2327244 to J.C.S., and the University of Texas at Austin, Graduate School Continuing Fellowship 2022-2023 to F.Y.

## Author contributions

F.Y., K.J.D., J.M.H., and J.C.S. designed the experiments. L.W. and E.M.L. purified Eps15 and the variants for *in vitro* experiments. F.Y., A.S. and B.T.M conducted the *in vitro* experiments. F.Y., S.G., and A.S. performed the cloning and cell assays. F.Y., S.G., G.A., and J.C.S. contributed to data analysis. F.Y., L.W., E.M.L., J.M.H., and J.C.S. wrote the paper. All authors consulted on manuscript preparation and editing.

## Data availability

All data supporting this work are available on request from corresponding author. CMEanalysis codes are available here: https://www.utsouthwestern.edu/labs/danuser/software/. No custom code was generated for this study.

